# Ecdysone regulates the *Drosophila* imaginal disc epithelial barrier, determining the duration of regeneration checkpoint delay

**DOI:** 10.1101/2020.07.16.207704

**Authors:** Danielle DaCrema, Rajan Bhandari, Faith Karanja, Ryunosuke Yano, Adrian Halme

## Abstract

Regeneration of *Drosophila* imaginal discs, larval precursors to adult tissues, produces a systemic response, a regeneration checkpoint that coordinates regenerative growth with developmental progression. This regeneration checkpoint is coordinated by the release of the relaxin-family peptide Dilp8 from regenerating tissues. Secreted Dilp8 protein can be detected within the imaginal disc lumen. The disc epithelium separates from the lumen from the larval hemolymph and the targets for Dilp8 activity in the brain and prothoracic gland. Here we demonstrate that the imaginal disc epithelial barrier limits Dilp8 signaling and checkpoint delay. We also observe that the wing imaginal disc barrier becomes more restrictive during development, becoming impermeable only at end of the final larval instar. This change in barrier permeability is driven by the steroid hormone ecdysone and correlates with changes in localization of Coracle, a component of the septate junctions that is required for the late, impermeable epithelial barrier. Based on these observations, we propose that the imaginal disc epithelial barrier regulates the duration of the regenerative checkpoint, providing a mechanism by which tissue function can signal the completion of regeneration.

**Summary Statement:** Ecdysone signaling directs the *Drosophila* third instar imaginal disc epithelial barrier to mature, becoming more restrictive. This mature barrier limits Dilp8 signaling and determines the duration of the regeneration checkpoint.

## Introduction

*Drosophila melanogaster* imaginal discs, larval precursors to adult organs, can regenerate after damage early in development, but lose this regenerative ability to prior to pupariation (Halme et al., 2010). Following damage, regenerating imaginal discs activate a regeneration checkpoint through release of the relaxin peptide *Drosophila* insulin-like peptide 8 (Dilp8) (Colombani et al., 2012; Garelli et al., 2012). Dilp8 functions in the brain and the prothoracic gland (PG) by binding to the relaxin receptor (Lgr3), which inhibits the synthesis of the steroid hormone ecdysone (Colombani et al., 2015; Garelli et al., 2015; Jaszczak et al., 2016; Vallejo et al., 2015). Ecdysone triggers many of the developmental events that characterize the end of larval development and pupariation, including regeneration restriction (Hackney and Cherbas, 2014; Halme et al., 2010). Therefore, the inhibition of ecdysone synthesis during the regeneration checkpoint extends the larval phase, coordinating regeneration with developmental progression (Halme et al., 2010; Jaszczak et al., 2016). However, it is unclear what events determine the duration of the regenerative checkpoint and signal the completion of regeneration, allowing the larvae to enter pupation and progress through development to the adult stage.

Colombani et al. observed Dilp8 in the luminal space of the wing imaginal discs, between the primary epithelia and the peripodial membrane (Colombani et al., 2012). Since the imaginal discs derive from the larval epidermis, emerging like an inflating balloon into the larval body cavity, this luminal space is topologically separated from the hemolymph by the imaginal disc epithelia (Pastor-Pareja et al., 2004). This led us to hypothesize that the imaginal disc epithelial barrier activity might regulate Dilp8 signaling by preventing access of Dilp8 in the disc lumen to the larval hemolymph, thus blocking signaling through the Lgr3 receptors in the brain and PG.

The epithelial barrier is a semi-permeable diffusion barrier between adjacent epithelial cells, and is formed by tight junctions in vertebrates and septate junctions in invertebrates (Tepass et al., 2001). Claudin proteins determine epithelial barrier exclusivity by homo- and heterodimerizing with the claudins of neighboring cells (Furuse and Tsukita, 2006). The claudins are localized to and stabilized at the junctions by a large core complex (Izumi and Furuse, 2014). However, the assembly and function of each member of the septate junction complex is not well understood. One subcomplex of the septate junction includes Coracle (Cora), a member of the Protein 4.1 superfamily (Fehon et al., 1994), and Neurexin-IV (Nrx) (Baumgartner et al., 1996). *In vivo*, Cora and Nrx both localize to the septate junctions and are necessary for stabilization of claudins at the septate junction and barrier activity (Baumgartner et al., 1996; Fehon et al., 1994; Genova and Fehon, 2003).

Here we describe experiments demonstrating that the epithelial barrier of wing imaginal discs matures during the third instar in response to increasing ecdysone levels and a re-localization of Coracle along the lateral membrane. This mature, prepupal epithelial barrier limits Dilp8 signaling and determines the duration of the developmental checkpoint after damage.

## Results

### The activity of the Dilp8 is constrained by the imaginal disc epithelial barrier, determining the duration of regeneration checkpoint delay

Previously, Colombani et al. observed that Dilp8 protein could be detected in the luminal space between the primary wing disc epithelia and the peripodial membrane, within a region of the imaginal disc that is topologically separate from the larval hemolymph (Colombani et al., 2012; Pastor-Pareja et al., 2004). We recapitulated these findings by exogenously expressing a FLAG epitope-tagged allele of Dilp8 (Garelli et al., 2012) in wing imaginal discs with *apterous-Gal4* (*Ap-Gal4*) to induce expression in the dorsal half of the tissue (Cohen et al., 1992). We detected an accumulation of Dilp8::FLAG in the lumen of the wing imaginal discs (Fig. S1). This observation led us to hypothesize that this luminal localization might limit Dilp8 signaling by preventing access to Lgr3 receptors the brain and prothoracic gland, targets of Dilp8 to regulate growth and developmental timing (Colombani et al., 2015; Garelli et al., 2015; Vallejo et al., 2015).

As the imaginal disc is an epithelial tissue, we examined whether disruption of the epithelial barrier would produce increased developmental checkpoint signaling when Dilp8 is expressed in the wing disc. To test whether the imaginal disc epithelial barrier constrains Dilp8 signaling we expressed Dilp8 in the wing imaginal disc and measured the effect on developmental checkpoint delay when co-expressing an RNAi construct against a necessary component of the imaginal disc epithelial barrier, the claudin Kune-kune (*kune*^*RNAi*^, see Fig. 2A,B and S8B,C for demonstrations of RNAi activity) (Nelson et al., 2010). We used *beadex-Gal4* (*Bx-Gal4*), to express either *dilp8::FLAG, kune*^*RNAi*^, or both constructs in the pouch region of the wing disc (Milán et al., 1998) and measured developmental checkpoint delay relative to *lacZ* expressing control larvae. While *Bx>dilp8::FLAG* and *Bx>kune*^*RNAi*^ expression each produce a short delay in development relative to control larvae (13 and 10 hours respectively), the co-expression of Dilp8::FLAG and *kune*^*RNAi*^, produces a strong genetic interaction and a synergistic effect on delay (40 hours; Fig. 1A). To confirm this synergistic effect on delay is not due to increased activity at the endogenous *dilp8* locus, we examined the effect of co-expression of Dilp8::FLAG and *kune*^*RNAi*^ in a homozygous *dilp8* hypomorphic genetic background. Even without functional endogenous copies of the *dilp8* gene, we still observe a strong genetic interaction between Dilp8::FLAG and *kune*^*RNAi*^ and synergistic effect on developmental checkpoint delay (Fig. 1B). We generated a similar synergistic interaction when we co-expressed Dilp8::FLAG and an RNAi construct targeting Neurexin IV (Nrx), another necessary component of the imaginal disc septate junction (Baumgartner et al., 1996) (Fig. S2). These data indicate that disrupting the epithelial barrier through either of two distinct genetic targets can produce synergistic extension of delay during Dilp8 expression. These results support the hypothesis that the epithelial barrier limits Dilp8 signaling from the wing imaginal disc by retaining Dilp8 in the wing disc lumen.

**Figure 1.**
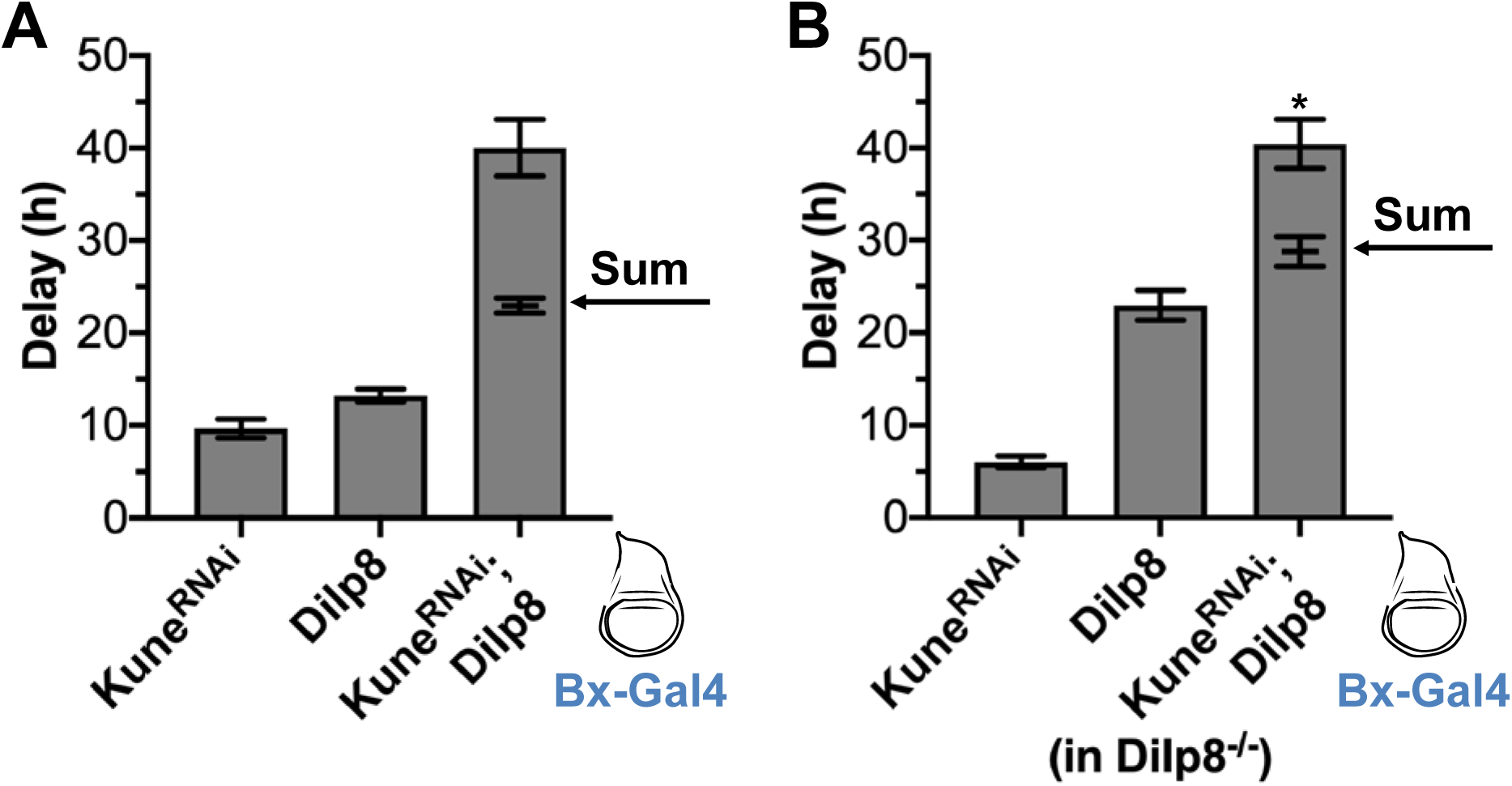
The epithelial barrier limits Dilp8 signaling. (A) Co-expression of *kune*^*RNAi*^ and Dilp8 induces synergistic delay. Ectopic expression of either *kune*^*RNAi*^, Dilp8, or co-expression of *kune*^*RNAi*^ and Dilp8 (*kune*^*RNAi*^; Dilp8) induce developmental delay compared to LacZ controls when expressed in the wing imaginal disc under Bx-Gal4 (expression region in blue). The delay induced by co-expression of *kune*^*RNAi*^ and Dilp8 (*kune*^*RNAi*^; Dilp8) is significantly more than the sum of the delay induced by *kune*^*RNAi*^ and Dilp8 expressed alone (sum indicated by arrow). (B) This trend holds true when endogenous Dilp8 is limited by expression in a Dilp8 hypomorphic background (Dilp8^MI00727^/Dilp8^MI00727^; Garelli et al., 2012). Data were collected from at least three independent experiments, bars represent mean ± SEM, * p < 0.05 from one sample t-test comparing the additive value and observed delay.

**Figure 2.**
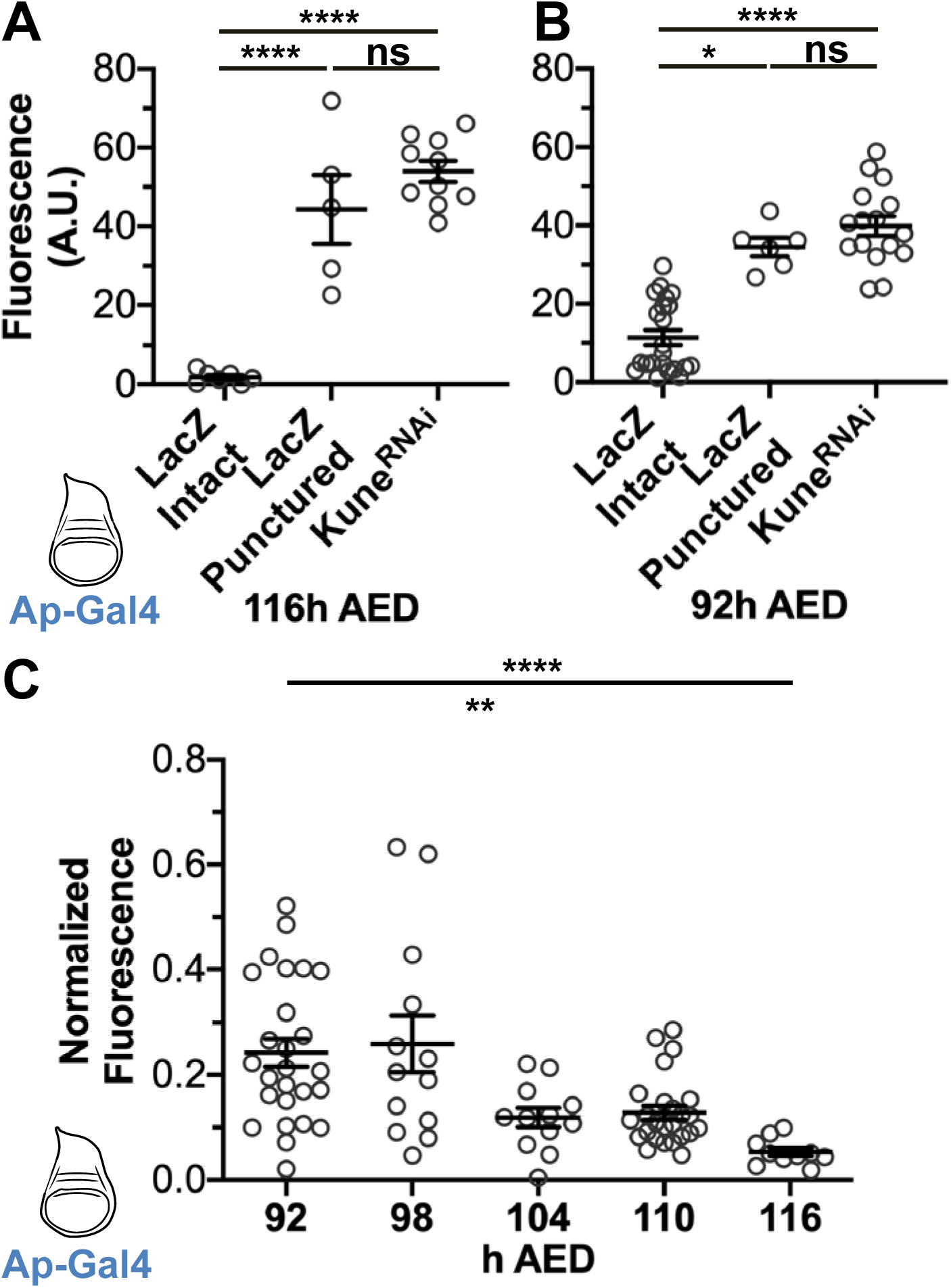
The wing disc epithelial barrier becomes more restrictive with progression through the third larval instar. (A-B) The wing imaginal disc epithelial barrier excludes 10 kD dextran and the function of the barrier is dependent on Kune at both (A) 116h and (B) 92h AED. LacZ and Kune^RNAi^ were expressed in the imaginal disc with Ap-Gal4 (expression area diagramed in blue). Imaginal discs were incubated in 10 kD fluorescein-conjugated dextran for 30 minutes prior to (see methods for details). Luminal intensity was measured in the LacZ controls, LacZ controls that were punctured during dissection prior to fixing, and the Kune^RNAi^ expressing discs. (C) The maturation of the epithelial barrier occurs gradually from 92h to 116h AED. Every 6 hours from 92h and 116h AED, barrier function was measured, as previously described, in wing imaginal discs expressing LacZ or Kune^RNAi^ with Ap-Gal4. Data indicate luminal intensity of intact LacZ expressing discs normalized to the mean luminal intensity of the Kune^RNAi^ expressing discs from the same timepoint, Kune^RNAi^ data represented in Figure S6. (A-C) Graphs represent mean ± SEM, with individual points indicating values of single images. Left to right, n = (A) 7, 5, 10, (B) 22, 6, 16, (C) 26, 13, 12, 25, 10. (A-C) ns not significant, * p < 0.05, ** p < 0.01, **** p < 0.0001 as calculated by (A,C) Brown-Forsythe and Welch ANOVA with Dunnett’s T3 test for multiple comparisons or (B) one-way ANOVA with Tukey test for multiple comparisons.

### The wing imaginal disc epithelial barrier becomes more restrictive during the last larval instar

The experiments above demonstrate that the wing disc epithelial barrier can limit Dilp8 signaling. However, Dilp8 expression in the wing disc produces developmental delay even when the wing disc epithelial barrier is not disrupted (Fig. 1A,B; Colombani et al., 2012; Garelli et al., 2012). Additionally, basal levels of Dilp8 expression in the developing wing disc regulate tissue symmetry through communication with Lgr3 in the brain (Colombani et al., 2012; Garelli et al., 2012; Garelli et al., 2015; Vallejo et al., 2015). To reconcile these observations with the sequestration of Dilp8 in the wing disc lumen during in late third instar larvae, we hypothesized that there may be changes in the epithelial barrier permeability during development. To examine this, we developed a quantitative method for measuring epithelial barrier permeability that is an extension of a method described by Lamb et al., in which the ability of fluorophore-conjugated dextran to enter into the lumen of a tissue was used to assess epithelial barrier function (Lamb et al., 1998). For our assay, we incubated inverted larval carcasses for 30 minutes in solution containing fluorescein-conjugated dextran prior to paraformaldehyde fixation. We mounted and imaged the fixed imaginal discs, then measured the fluorescein signal in the lumen of the imaginal discs to quantify epithelial barrier permeability (see Materials and Methods and Fig. S3 for a more complete description of this assay). When we examined late third instar imaginal discs (116 hours After Egg Deposition; h AED), we observed that very little fluorescent signal is detected in the wing disc lumen (Fig. 2A). To determine whether exclusion of dextran from late larval wing discs is dependent on epithelial barrier function, we measured fluorescence in imaginal discs that had been punctured with a forceps tip prior to incubation with fluorescently-labeled dextran. As expected, we observed that puncturing the wing disc led to a substantial increase in luminal fluorescence detected in these discs after fixation (Fig. 2A), demonstrating that an intact epithelium is necessary for the exclusion of dextran from the lumen of late third instar wing discs. We then tested whether this exclusion reflected the activity of the epithelial barrier by measuring dextran infiltration into wing discs expressing *kune*^*RNAi*^ (*Ap>kune*^*RNAi*^). Consistent with a critical role for the claudin Kune in wing epithelial barrier function, we observed an equivalent amount of fluorescence infiltration into 116h AED *Ap>kune*^*RNAi*^ discs as we observed in punctured discs (Fig. 2A). Therefore, loss of *kune* appears to completely disrupt the epithelial barrier in late third instar wing discs.

To characterize the epithelial barrier exists in earlier wing discs, we examined wing discs 24 hours earlier in development (92h AED, about the middle of the third and last larval instar), a time when disc damage and/or Dilp8 expression is still capable of producing developmental delay. In 92h AED discs we observe that physical puncture and *kune*^*RNAi*^ expression both produce increases in dextran infiltration into the wing disc lumen (Fig. 2B), similar to what we observed at 116h AED. This indicates, that wing imaginal discs in the middle of the third instar have a functioning epithelial barrier mediated by *kune*. However, we also noticed that the level of dextran infiltration into control 92h AED wing discs appeared to be much higher than the fluorescence observed in the same tissues at 116h AED (compare Fig. 2A and B), suggesting that the epithelial barrier of the wing disc grows more restrictive as the larvae approach the end of the third instar. This change in barrier permeability from 92h to 116h AED is also observed with 70 kD dextran (Fig. S4), suggesting that the change in permeability does not reflect a change in size selectivity of the barrier. To further characterize the maturation of the more restrictive barrier, we used our quantitative barrier permeability assay to examine dextran infiltration at six-hour intervals between 92h AED and 116h AED. We normalized the fluorescence intensity to discs of equivalently staged larvae with barriers disrupted by *kune*^*RNAi*^ expression. Consistent with our earlier observations at 92h and 116h AED, as the wing disc develops, we see a progressive decrease in barrier permeability that limits the infiltration of dextran into the lumen of this tissue (Fig. 2C, for an illustration of the individual fluorescence distributions, see Fig. S5).

Based on these observations, we conclude that the wing disc epithelial barrier limits diffusion throughout the third instar since we see an increase in permeability in punctured or *kune*^*RNAi*^ expressing discs at earlier stages. However, we see a substantial difference in permeability of the barrier between earlier and later discs, with the wing disc epithelial barrier becoming progressively less permeable as larvae advance through the third instar.

### Changes in epithelial barrier permeability correlate with changes in junctional protein expression and localization

The changes that we observe in barrier permeability during the last larval instar could be due to changes in localization and/or activity of different components of the epithelial barrier. To address this, we examined the localization of three major components of the epithelial barrier: the claudin Kune, which forms the intercellular component of the junctional barrier (Nelson et al., 2010), along with Cora and Nrx which together regulate the function of pleated septate junctions in ectoderm-derived tissues such as the imaginal discs (Baumgartner et al., 1996; Lamb et al., 1998). Using indirect immunofluorescent staining with a Kune protein-directed antibody (Nelson et al., 2010), we observed that Kune protein is localized to the apical region of the lateral membrane throughout the third instar (Fig. 3D,E; Fig. S6D,E), which is the region of septate junctions localization in imaginal tissues (Lamb et al., 1998; Ward IV et al., 2001). We also observed that there is a substantial increase in localized Kune signal at the septate junction in late third instar discs (116h AED) when compared to earlier (92h AED) discs (Fig. 3F, quantification method described in Materials and Methods and diagramed in Fig. S7). To localize Nrx, we used a functional Nrx-GFP fusion (Buszczak et al., 2007; Morin et al., 2001). Like Kune, Nrx-GFP also localizes to the septate junctions throughout the third instar and increases in signal intensity from 92 to 116 hAED (Fig. 3G-I; Fig. S6F,G). In contrast to Kune and Nrx-GFP, the localization of Cora is more dynamic. At 92h AED, Cora is localized either almost uniformly along the entire length lateral membrane, without selectivity for the septate junction, or with slight accumulation at the septate junctions (Fig. 3J,L; Fig. S6H). However, by 116h AED, Cora localization shifts to become restricted to an apical-lateral localization at the septate junctions where we observe Kune and Nrx-GFP signals (Fig. 3K,L; Fig. S6I).

**Figure 3.**
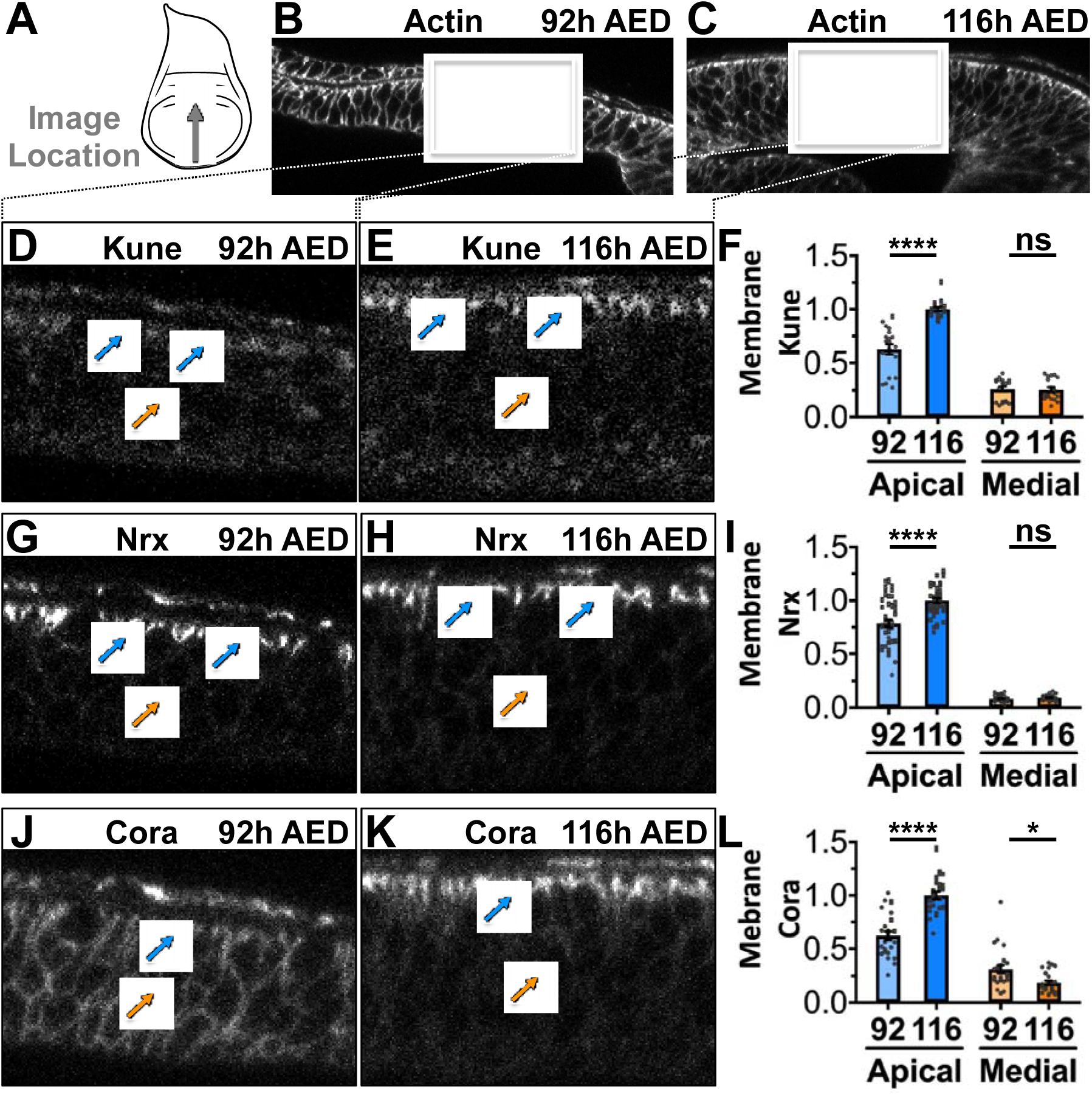
The localization of septate junction components changes between 92h and 116h AED. (A) Cross sections of the pouch region of the wing imaginal discs were collected by capturing X-Z images. All images were similarly oriented with the ventral portion of the disc facing left and the dorsal portion of the disc facing right. Approximate region and direction of the images is indicated. (B-C) Disc area was determined by Actin (rhodamine phalloidin) staining at (B) 92h and (C) 116h AED. The box represents the area of focus for images in D, G and J for the 92h AED disc and E, H, and K for the 116h AED disc. Blue arrows indicate apical-lateral localization, orange arrows indicate medial-lateral localization (or lack thereof). (D-E) Representative images of Kune localization at (D) 92h and (E) 116h AED showing of apical-lateral Kune localization. (F) Quantification of Kune localized along the apical-lateral and medial-lateral membrane, normalized to mean Kune membrane intensity at 116h AED. (G-H) Representative images of Nrx localization at 92h (G) and 116h (H) AED showing apical-lateral localization. (I) Quantification of Nrx localized along the apical-lateral and medial-lateral membrane, normalized to mean Nrx membrane intensity at 116h AED. (J-K) Representative images of Cora localization at (J) 92h and (K) 116h AED showing diffuse Cora localization at 92h AED, and apical-lateral Cora localization at 116h AED. (L) Quantification of Cora localized along the apical-lateral and medial-lateral membrane, normalized to mean Cora membrane intensity at 116h AED. (D,E,G,H,J,K) Full images in Figure S6. (F,I,L) Individual points represent mean ± SEM for each image, with the n considered to be the number of cell-cell interactions across the region measured. Bars represent mean ± SEM across the images, with the n considered to be the number of images measured. Details about the quantification method are explained in the Methods Section and Figure S7. n = (F) 19, 92h AED and 18, 116h AED images, (I) 39, 92h AED and 41, 116h AED images, and (L) 24, 92h AED and 24, 116h AED images. ns not significant, * p < 0.05, **** p < 0.0001 as calculated by unpaired, two-tailed t-test, except for apical-lateral Kune, which had unequal variance, in this case significance was calculated by unpaired, two-tailed t-test with Welch’s correction.

In summary, we observe changes in core SJ localization that correlate with the decrease in permeability of the epithelial barrier: Increased Kune localization at the septate junction, and a refinement of Cora localization, from a uniform distribution along the lateral membrane to localization at the apical site of the septate junction (Fig. 5G).

**Figure 4.**
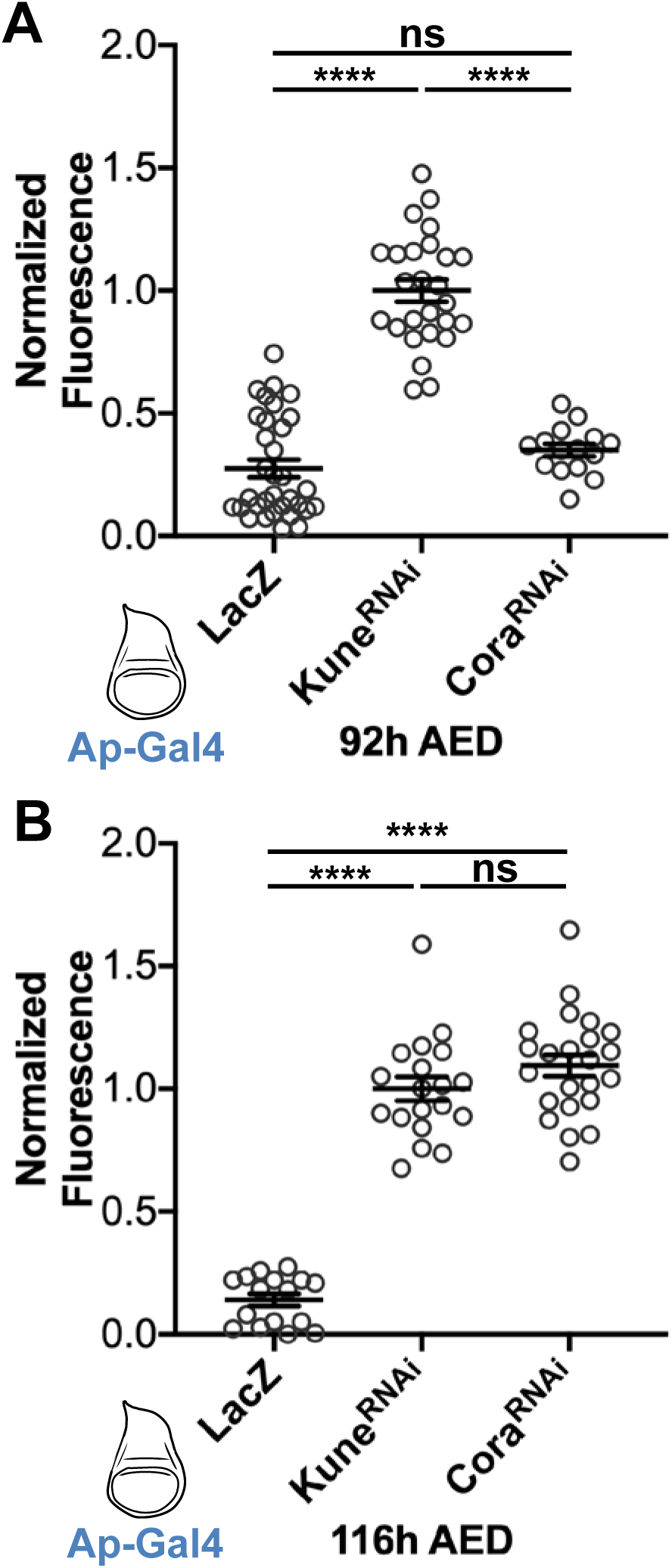
The role of Cora in epithelial barrier activity changes from 92h and 116h AED. (A-B) Function of the epithelial barrier in discs expressing *lacZ, kune*^*RNAi*^, or *cora*^*RNAi*^ (using Ap-Gal4, expression diagramed in blue) to exclude 10 kD dextran at (A) 92h and (B) 116h AED. At 92h AED, the barrier of *cora*^*RNAi*^ expressing discs is similar to *lacZ* expressing discs. At 116h AED, the barrier of *cora*^*RNAi*^ expressing discs is similar to *kune*^*RNAi*^ expressing discs. Data are normalized to the mean luminal intensity of *kune*^*RNAi*^ expressing discs. Graph represents mean ± SEM, with individual points indicating values of single images. Left to right, n = (A) 33, 26, and 15 images, and (B) 16, 19, and 23 images. ns not significant, **** p < 0.0001 as calculated by Brown-Forsythe and Welch ANOVA with Dunnett’s T3 test for multiple comparisons.

**Figure 5.**
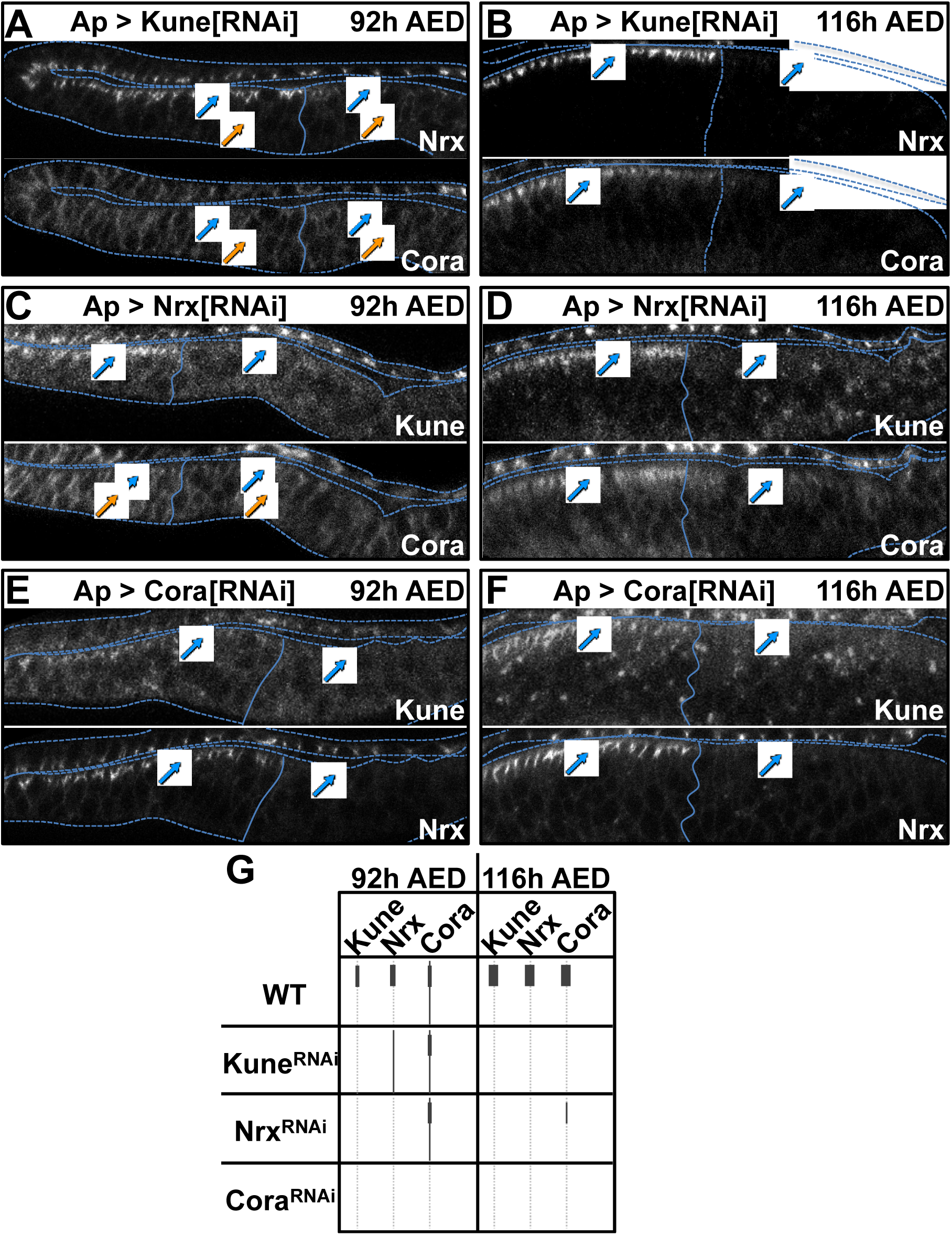
The localization of Nrx and Kune is dependent on Cora at both 92h and 116h AED. Ap-Gal4 was used to express *kune*^*RNAi*^, *nrx*^*RNAi*^, or *cora*^*RNAi*^ and the localization of the Kune, Nrx, and Cora were assessed (images with same protein as the RNAi are in Figure S8). Images were taken spanning the dorsal-ventral boundary (solid line, defined by data in Figure S8) and oriented with the dorsal region (expression area) on the right. Dotted lines represent tissue outline defined by Actin staining (Figure S8). Blue arrows indicate apical-lateral localization, orange arrows indicate medial-lateral localization. (A-B) Localization of Nrx and Cora in *Ap > kune*^*RNAi*^ expressing discs at (A) 92h and (B) 116h AED. In the portion of the disc without *kune*^*RNAi*^ expression Nrx is apical-laterally localized at 92h and 116h AED, without medial-lateral localization. Nrx intensity localization is lost with *kune*^*RNAi*^ expression at both times, but at 92h AED, also becomes more diffusely localized. Cora localization is independent of Kune at 92h AED, but is lost with *kune*^*RNAi*^ expression at 116h AED. (C-D) Localization of Kune and Cora in *Ap > nrx*^*RNAi*^ expressing discs at (C) 92h and (D) 116h AED. Kune localization is lost with *nrx*^*RNAi*^ expression at both times. Cora localization is independent of Kune at 92h AED, but is lost with *kune*^*RNAi*^ expression at 116h AED. Cora localization is independent of Kune at 92h AED, but is significantly diminished, although not completely lost, with *kune*^*RNAi*^ expression at 116h AED. (E-F) Localization of Kune and Nrx in *Ap > cora*^*RNAi*^ expressing discs at (E) 92h and (F) 116h AED. Kune and Nrx localization are lost with *cora*^*RNAi*^ at both times. (G) Summary of imaging data, dotted line represents a cell membrane. (A-F) Images are representative from n = (A) 8, (B) 10, (C) 8, (D) 6, (E) 5, and (F) 11 images.

### Coracle is required to produce the changes in epithelial barrier permeability during the last larval instar

To determine whether Cora activity is important for the observed decrease in septate junction permeability during the last larval instar, we examined the effect of *cora*^*RNAi*^ expression on wing imaginal disc barrier permeability. At 92h AED *Ap>cora*^*RNAi*^ expression has little impact on the wing epithelial barrier permeability. The barrier activity in Cora knockdown discs is similar to control discs, with much greater selectivity than *Ap>kune*^*RNAi*^ expressing discs, which have no functioning barrier (Fig. 4A). In contrast, in late third instar wing imaginal discs (116h AED) *Ap>cora*^*RNAi*^ expressing discs exhibit a completely disrupted epithelial barrier, comparable to that seen in *Ap>kune*^*RNAi*^ expressing discs (Fig. 4B). Therefore, as the third instar progresses, there is a change in the role of Cora for septate junction barrier activity: 92h AED wing discs have a weaker, somewhat permeable, barrier activity that requires Kune, but does not depend on Cora, whereas 116h AED wing discs have a more restrictive barrier activity that is completely dependent on Cora.

Cora, Kune, and Nrx localize interdependently at the septate junctions during the development of the embryonic tracheal epithelia (Nelson et al., 2010; Oshima and Fehon, 2011). We examined whether the same interdependence occurs in the wing imaginal disc, and whether it changes during development. To do this, we visualized the localization of Cora, Kune, and Nrx-GFP in discs in which we had knocked down each of these components using RNAi targeting constructs driven by *Ap-Gal4*. Predictably, when we expressed RNAi lines targeting Cora, Kune, and Nrx, we saw a loss of expression of the targeted gene product in the dorsal compartment of both 92 and 116h AED wing discs (Fig. S8), demonstrating the efficacy of the RNAi constructs. We then examined the interdependence for localization of these three septate junction proteins at 92h and 116h AED wing discs (Fig. 5 A-F, summarized in Fig. 5G). At 92h AED, the localization of Cora along the lateral membrane is unaffected by RNAi targeted knockdown of either Kune (*Ap>kune*^*RNAi*^, Fig. 5A) or Nrx (*Ap>nrx*^*RNAi*^, Fig. 5C). However, the localization of Cora at the septate junctions at 116h AED is disrupted by RNAi targeted knockdown of either Kune (*Ap>kune*^*RNAi*^, Fig. 5B) or Nrx (*Ap>nrx*^*RNAi*^, Fig. 5D). Therefore, the early, diffuse localization of Cora along the lateral membrane does not require the activity of either Kune or Nrx, but its refinement to the septate junction at the end of the larval period depends on both Kune and Nrx. Nrx localization at the septate junctions in 92h AED larvae is dependent on both Kune (*Ap>kune*^*RNAi*^, Fig. 5A) and Cora (*Ap>cora*^*RNAi*^, Fig. 5E). However, in the absence of Kune, we see faint Nrx localization along the lateral membrane (Fig. 5A, orange arrow), which might reflect an association with Cora, whereas in the absence of Cora, we see less evidence of lateral localization of Nrx (Fig. 5E, arrowhead). At 116h AED, Nrx localization at the septate junction is entirely dependent on both Kune and Cora (Fig. 5B,F) and no lateral redistribution of Nrx is seen in either of these mutants. Finally, the septate junction localization of Kune at both 92 and 116h AED is entirely dependent on Nrx (*Ap>nrx*^*RNAi*^, Fig. 5C,D) and, surprisingly, Cora (*Ap>cora*^*RNAi*^, 5E,F). The requirement for Cora to localize Nrx and Kune at 92h AED is unexpected since Cora is not required for the barrier activity observed at 92h AED (Fig. 4A). We address some possible explanations of this observation in the Discussion section.

As summarized in Figure 5G, we see a complex interdependence between Kune, Nrx, and Cora that determines their localization as the wing disc epithelial barrier becomes more restrictive during the larval third instar. These data indicate that the epithelial barrier becomes more mature and restrictive, barrier activity becomes dependent on Cora as it re-localizes to the septate junctions.

### Ecdysone signaling promotes decreased epithelial barrier permeability and Coracle re-localization

The steroid hormone ecdysone is a critical endocrine regulator of *Drosophila* developmental progression. During the third larval instar, pulses of ecdysone synthesis drive a progressive increase in ecdysone titer throughout the larva that promotes growth and differentiation of the imaginal discs (Colombani *et al.* 2005; Lavrynenko *et al.* 2015. During the activation of the regenerative checkpoint following imaginal disc damage, Dilp8 release suppresses ecdysone synthesis through Lgr3 receptors in both the larval brain and PG (Hackney et al., 2012; Halme et al., 2010; Jaszczak et al., 2016).

To assess whether the changes we observe in wing disc epithelial barrier permeability are driven by ecdysone signaling, we first tested whether increasing ecdysone titer in larvae would reduce disc barrier permeability. To do this, we transferred 80h larvae to either food containing 0.6 mg/ml 20-hydroxyecdysone dissolved in ethanol, or food with ethanol alone as a control and assessed wing disc barrier function at 98h AED using our dextran infiltration assay (illustrated Fig. S9A). In previously published work we have seen that this concentration of 20-hydroxyecdysone can alter ecdysone titer, but doesn’t substantially accelerate pupariation timing (Colombani et al., 2005; Jaszczak et al., 2015). When we assessed barrier activity in ecdysone-fed larvae, we see that wing disc permeability is substantially reduced when compared with wing discs from control larvae (Fig. 6A; Fig. S9B), similar to what we observe in more mature (116h AED) wing discs (compare to Fig. 1C). This result indicates that increasing ecdysone titers can promote the development of the restrictive barrier we see as the third larval instar progresses.

**Figure 6.**
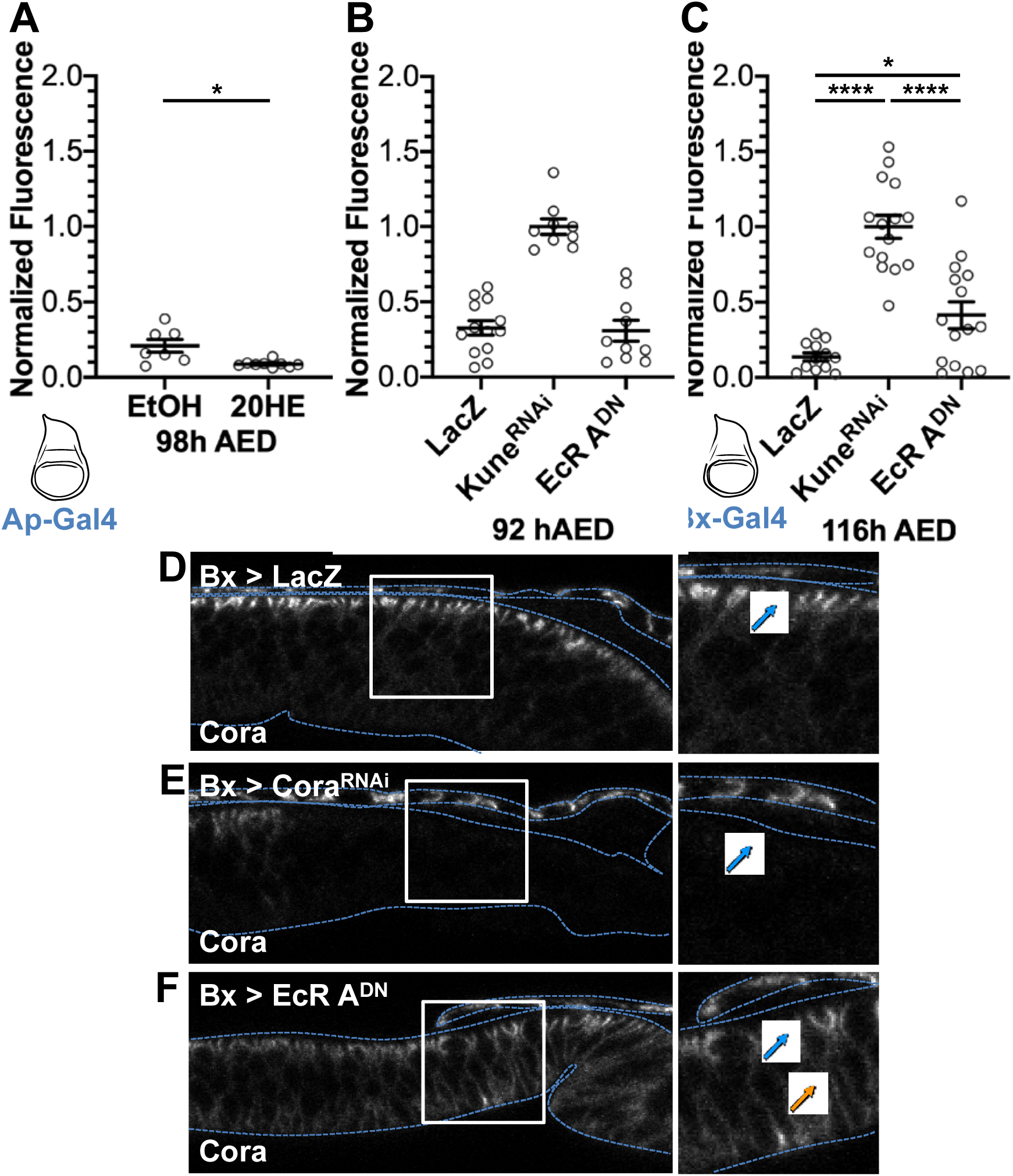
Ecdysone induces barrier maturation and Cora localization. (A) Function of the epithelial barrier at 98h AED in the wing imaginal discs of larvae that were switched to food containing ethanol control (EtOH) or 0.6 mg/mL 20-hydroxecdysone (20HE) at 80h AED. Barrier function is normalized to *Ap > kune*^*RNAi*^ expressing discs under the same feeding conditions (complete data in Figure S9B). Expression area diagramed in blue. (B-C) Epithelial barrier function of *Bx > lacZ* (wild type control), *Bx > kune*^*RNAi*^, and *Bx > EcR A*^*DN*^ at (B) 92h and (C) 116h AED. Barrier function is normalized to *Bx > kune*^*RNAi*^ expressing discs under the same feeding conditions. Expression area diagramed in blue. (D-F) Localization of Cora in (D) *Bx > lacZ* (wild type control), (E) *Bx > kune*^*RNAi*^, and (F) *Bx > EcR A*^*DN*^ at 116h AED. Dotted lines indicate tissue outline as defined by actin staining (rhodamine phalloidin; Figure S10). Tissues are oriented with the dorsal region of the pouch (higher level of Bx expression) on the right. White box indicates zoomed area to the right. Blue arrows indicate apical-lateral Cora localization (or lack thereof in *cora*^*RNAi*^), orange arrow indicates medial-lateral localization. (A-C) Graphs represent mean ± SEM, with individual points indicating values of single images. Left to right, n = (A) 7 and 9, (B) 13, 8, 10, and (C) 12, 15, 15. (A-C) ns not significant, * p < 0.05, **** p < 0.0001 as calculated by (A) Unpaired t-test with Welch’s correction, (B) Ordinary one-way ANOVA with Tukey’s multiple comparisons test, or (C) Brown-Forsythe and Welch ANOVA tests with Dunnett’s T3 multiple comparisons test. (D-F) Images are representative of n = (D) 7, (E) 7, and (F) 5 images.

To determine whether ecdysone acts directly on the wing disc and is necessary for the change in barrier permeability, we assessed barrier function in wing discs expressing the dominant-negative ecdysone receptor allele *EcR.A*^*W650A*^ in wing discs using *Bx-Gal4*. In 92h AED discs, limiting ecdysone signaling produces little effect on barrier permeability (Fig. 6B), demonstrating that the barrier function doesn’t rely on ecdysone signaling at this earlier stage. However, in 116h AED discs we see that blocking ecdysone signaling now produces a substantial increase in epithelial barrier permeability (Fig. 6C). The expression of *EcR.A*^*W650A*^ at 116h AED doesn’t produce the same disruption in barrier function as expression of *kune*^*RNAi*^, rather it produces barrier permeability similar to that seen in 92h AED discs (compare Figs. 6B and 6C). These results demonstrate that ecdysone signaling in the wing disc is not necessary for barrier activity in 92h AED wing discs, but is required for the maturation of the more restrictive barrier during the third instar.

Since a re-localization of Cora from the lateral membrane to the apical site of the septate junction is associated with the change in barrier permeability, we wanted to determine whether this change in Cora localization depends on ecdysone signaling in the wing disc. To do this, we examined the cellular localization of Cora in 116h AED control and *EcR.A*^*W650A*^ wing discs. Inhibition of ecdysone signaling in the wing disc produces a redistribution of Cora, from tightly localized to the apical lateral membrane, at the site of the septate junction, to a more uniform distribution along the lateral membrane (compare Fig. 6D and Fig. 6F; Actin staining Fig. S10), similar to what is observed in earlier 92h AED discs (Fig. 3D). This re-localization of Cora is consistent with how ecdysone effects the epithelial barrier, producing an increase in permeability that is similar to what is seen in 92h AED wing discs, but not completely disrupting the epithelial barrier (Fig. 5A).

In summary, we see that the epithelial barrier activity of the wing disc at 92h AED is not dependent on ecdysone signaling, whereas the re-localization of Coracle to the site of the septate junction, along with the decreased permeability of the epithelial barrier, are both dependent on ecdysone signaling within the wing imaginal disc epithelium.

### The epithelial barrier regulates the end point of regeneration

Our data indicate that ecdysone regulates a maturation of the epithelial barrier and that the epithelial barrier limits Dilp8 signaling. This led us to question whether the epithelial barrier determines the duration of the regenerative checkpoint. To test this, we targeted damage to the wing discs by using *Bx-Gal4* to express the TNFα homologue Eiger in the pouch of the wing disc (Igaki et al., 2002; Kauppila et al., 2003; Moreno et al., 2002), and examined the effects of barrier disruption on checkpoint duration. As previously observed, *kune*^*RNAi*^ expression alone produces only a minor effect on delay, whereas Eiger expression produces a substantial delay of 57 hrs. When we combine Eiger and *kune*^*RNAi*^ expression, we see a synergistic effect on delay that is significantly longer than the expected additive effect of *kune*^*RNAi*^ and Eiger expression alone (80 hours actual, 62 hours additive; Fig. 7A). However, this additional delay is not due to increased Dilp8 expression (Fig. S12), consistent with the epithelial barrier limiting Dilp8 signaling at the end of the regenerative checkpoint. We observed a similar result when the epithelial barrier is disrupted with *nrx*^*RNAi*^ (Fig. S11; Fig. S12).

**Figure 7.**
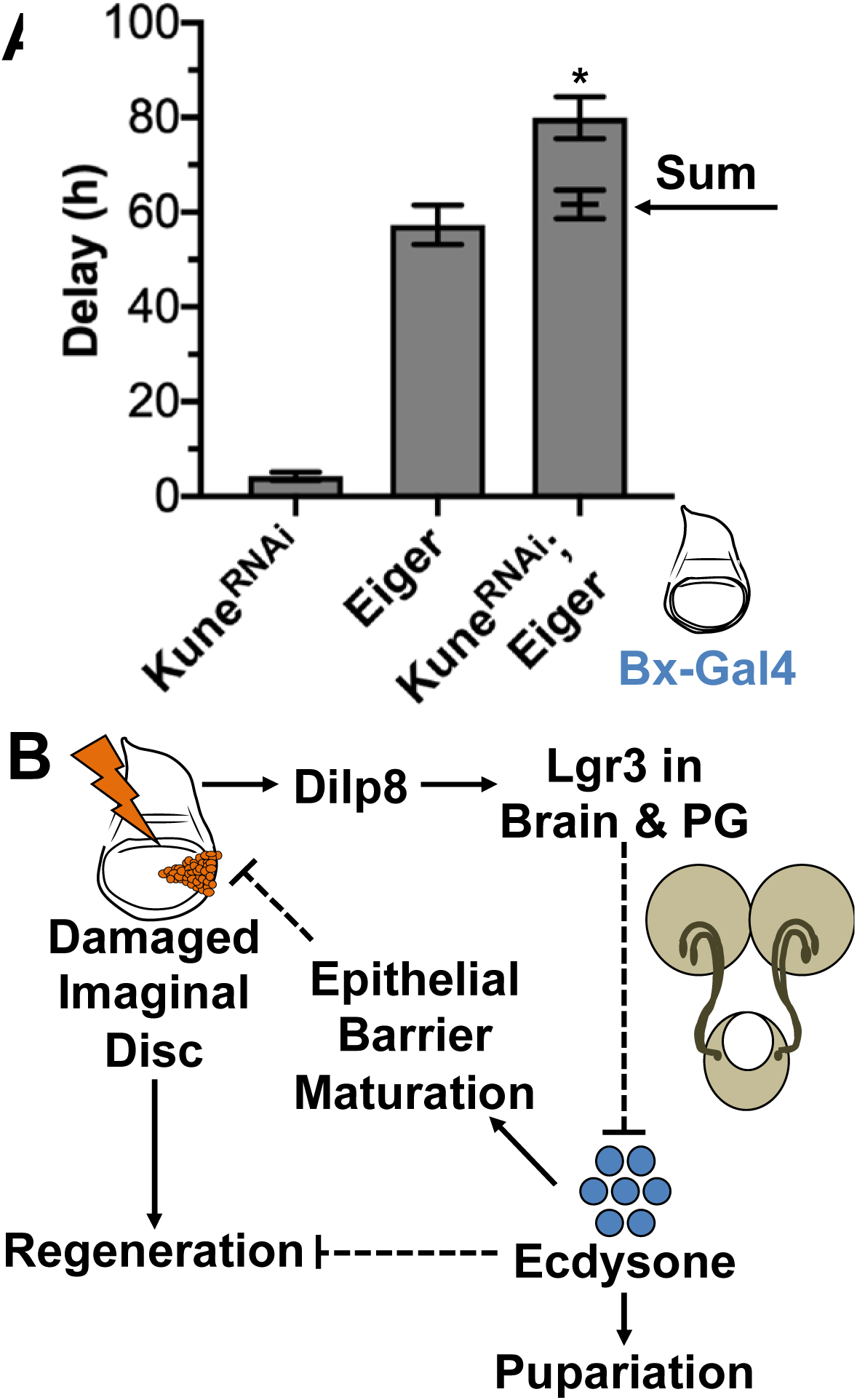
The epithelial barrier regulates the end point of regeneration. (A) Expression of *kune*^*RNAi*^ in eiger-damaged tissues induces synergistic delay. Data were collected from at least three independent experiments, bars represent mean ± SEM, * p < 0.05 from one sample t-test comparing the additive value and observed delay. (B) Model. Following damage to the imaginal discs, regeneration is initiated and Dilp8 is produced in the damaged tissues and is secreted. Dilp8 functions on Lgr3 receptors in the brain and PG to inhibit ecdysone production, resulting in a delay to pupariation. In the late third instar, high levels of ecdysone inhibit regenerative ability and also induce the maturation of the epithelial barrier in imaginal discs. The epithelial barrier inhibits Dilp8 signaling and regulates the duration of the regeneration checkpoint in development. We hypothesize that the restoration of the barrier following regeneration begins to limit Dilp8 release from the wing disc, and that this is the rate-limiting step for ending the regeneration checkpoint.

Together, these results demonstrate that a fully-functional epithelial barrier limits the duration of damage-induced checkpoint delay. Likely through the sequestration of Dilp8 within the imaginal disc lumen.

## Discussion

### How is the end of regeneration determined?

Despite progress towards understanding the cues that initiate regeneration and the signaling pathways that contribute to regenerative growth and repatterning, the mechanisms for determining when the target of regeneration is reached remain poorly understood (Fox et al., 2020). Here, we demonstrate that the formation of a functional, mature epithelial barrier determines the duration of the regenerative period by regulating Dilp8 signaling. Since Dilp8 is seen in the imaginal disc lumen (Colombani et al., 2012; Fig. S1), it seems likely that the epithelial barrier limits Dilp8 signaling by physically sequestering Dilp8 protein in the lumen of the imaginal disc, separated from the hemolymph and access to the prothoracic gland and the brain, where Dilp8 acts through Lgr3 to inhibit ecdysone production (Colombani et al., 2012; Colombani et al., 2015; Garelli et al., 2012; Garelli et al., 2015; Jaszczak et al., 2016). Therefore, we propose that as regeneration is completed, the balance between Dilp8 signaling (which delays developmental progression and limits ecdysone synthesis) and ecdysone signaling (which shuts down regenerative activity and promotes the advancement from the larval to the pupal phase of development) is shifted to favor ecdysone signaling by the establishment of the mature, prepupal epithelial barrier which traps Dilp8 in the regenerated disc lumen. In this way, the re-establishment of a restrictive epithelial barrier would be one mechanism for epithelial tissues to communicate their functional restoration and the completion of regeneration (model, Fig. 7B).

### Regulated maturation of the epithelial barrier

In these experiments, we also demonstrate that the function of the epithelial barrier changes during the third instar, growing more restrictive prior to pupariation in response to ecdysone. Although we were unable to determine the direct target of EcR that produces this change in barrier activity, we observe that ecdysone triggers the re-localization of Cora from a pattern of diffuse localization along the length of the lateral membrane to a specific localization at the septate junctions in the apical lateral membrane. This re-localization correlates with the establishment of a mature, restrictive epithelial barrier. This mechanism is similar to observations that Cora re-localizes to the septate junctions during the embryonic stages 12 and 17 in the developing salivary gland and embryonic epidermis, which is when the epithelial barrier is established in these tissues (Hall and Ward, 2016; Oshima and Fehon, 2011; Paul et al., 2003). Similar to the end of larval development, there is a peak of ecdysone production during this embryonic period and EcR is expressed in both these maturing epithelia (Kozlova and Thummel, 2000; Kozlova and Thummel, 2003; Tan et al., 2014). This leads us to suggest that the regulation of Cora localization by ecdysone may be a general mechanism for the maturation of a restrictive barrier in developing *Drosophila* epithelia.

However, our examination of epithelia barrier maturation in the wing disc also raises some interesting questions about the role of individual septate junction components which remain to be answered. In particular, the role is of the core components Cora and Kune in barrier function earlier in the third instar, at 92h AED. The barrier at this time is not as restrictive as is observed later, but still limits the passage of 10kD dextran molecules (Fig. 2B, compare Intact to Punctured). Kune is required for this early barrier activity but Cora is not necessary (Fig. 4). However, when we block Cora expression with an RNAi, we see that Cora is necessary for the localization of both Nrx and Kune at both 92h and 116h AED (Fig. 5). This suggests that in Cora mutant tissues at 92h AED, the mis-localized Kune (and Nrx) still retain some residual barrier activity, which is lost at 116h AED. One possible explanation is that a low level of Nrx and Kune localization along the lateral membrane may remain in Cora mutant tissues, but was undetectable by our imaging. This low level of Nrx and Kune localization may be sufficient to support early barrier activity in early discs, but not a similar barrier activity at the end of the third instar. Further study will be necessary to better understand how each of these core components contribute to the barrier as the wing disc matures.

### Could other signaling pathways be regulated by epithelial barrier maturation?

Our observation that Dilp8 is constrained by the wing disc epithelial barrier raises the question of whether other signals are regulated by sequestration in the imaginal disc lumen. One interesting possibility is the morphogen Decapentaplegic (Dpp, *Drosophila* BMP2/4 ortholog). Dpp has numerous, critical roles in growth and patterning across development, including in imaginal discs (Hamaratoglu et al., 2014). Setiawan et al. showed that during larval development, Dpp produced in the imaginal tissues inhibits ecdysone production in the prothoracic gland early in the third larval instar. By the late third instar, Dpp can no longer be detected in the larval hemolymph and the Dpp activity in the prothoracic gland ceases, despite high levels of Dpp expression in imaginal discs. Setiawan et al. hypothesized that this was due in part to the dilution of circulating Dpp as a result of increased hemolymph volume and the trapping of Dpp in imaginal disc tissues, but were unable to identify how the tissues trapped Dpp (Setiawan et al., 2018). Our data suggest that Dpp might be trapped in late third instar discs by the maturing epithelial barrier. Further experiments will be necessary to test this hypothesis.

In summary, our data demonstrate that in *Drosophila*, ecdysone signaling alters the permeability of the wing disc epithelial barrier during the third and final larval instar. We also show that mature, restrictive wing disc epithelial barrier limits Dilp8 signaling, determining the duration of the regenerative checkpoint. This provides an interesting mechanism by which the barrier, as a primary characteristic of epithelial tissues, can report the successful completion of regeneration.

## Materials and Methods

### Drosophila stocks and husbandry

The fly stocks used were, or were generated from crosses with, Ap-Gal4; UAS-LacZ.NZ, UAS-Dcr2/ SM6-TM6B (derived from Bloomington 3041), Bx-Gal4, UAS-Dcr2 (Dr. David Bilder), UAS-LacZ.NZ (BL3955), UAS-Kune[RNAi] (VDRC GD3962), UAS-NrxIV[RNAi] (VDRC GD8353), UAS-Sinu[RNAi] (VDRC GD44928), UAS-Cora[RNAi] (Bloomington 51845), UAS-EcR.A.W650A (Bloomington 9451), NrxIV-GFP (Bloomington 50798), UAS-eiger, UAS-Dilp8::3xFLAG (Dr. Maria Dominguez) (Garelli et al., 2012), and Dilp8::GFP (Bloomington 33079).

Stocks and crosses were maintained in 25° incubators with a 12-hour alternating light-dark cycle. Developmental timing was synchronized through egg staging, with collection from a 4-hour egg-laying interval on grape agar plates (Genesee Scientific) with a small amount of baker’s yeast paste. At 24h AED, 20-30 first instar larvae were transferred into vials containing cornmeal-yeast-molasses media (Archon Scientific B101). Ecdysone food was prepared by dissolving 1.2 mg 20-hydroxyecdysone (Sigma) dissolved in 95% ethanol in 2 mL of food media (final concentration 0.6 mg/mL), or an equivalent volume of ethanol for control. Larvae were reared as previously described until 80h AED then transferred to the ecdysone or ethanol-control food, approximately 6 larvae per vial (Halme et al., 2010).

For specific genotypes used in each Figure see Supplementary Materials and Methods.

### Dissection and Immunofluorescent Staining

Larvae were inverted and cleaned in PBS then fixed with 4% paraformaldehyde in PBS (20 min) and washed with PBS (twice for 5 min each). The tissues were permeabilized with 0.3% Triton in PBS (twice for 10 min each) then washed with a blocking solution of 10% goat and 0.1% Triton in PBS (30 min). Then the tissues were incubated rocking in primary antibody solutions (overnight at 4° or for two-to-four hours at room temperature). The process was repeated for secondary antibodies and then the tissues were incubated rocking in 80% glycerol in PBS (overnight at 4°). The tissues were stored at 4° in 80% glycerol and were mounted for imaging within one week of staining. Imaginal discs were isolated from the stained tissues and mounted on glass slides with Vectashield (Vector Laboratories). Cross-section images were taken from tissues mounted on slides with the coverslips raised by double-sided tape. For experiments with Kune or FLAG staining, the above procedure modified to reduce non-specific staining. In these experiments, larvae were dissected in Schneider’s Insect Medium (Sigma-Aldrich), fixed in 4% paraformaldehyde in Schneider’s Insect Medium (Sigma-Aldrich), stained in one day within three days of dissection, and imaged within three days of staining.

Antibody solutions were prepared in 10% goat serum and 0.1% Trion in PBS. The primary antibodies that were used are mouse β-Gal (1:250; Promega), mouse anti-Cora C615.16 (1:400; Developmental Studies Hybridoma Bank), mouse anti-FLAG M2 F1804 (1:250; Sigma-Aldrich), rabbit β-Gal (1:400; MP), and rabbit anti-Kune (1:1000; Dr. Mikio Furuse; Nelson et al., 2010). The secondary antibodies that were used are goat anti-mouse or anti-rabbit Alexa405, Alexa488, or Alexa633 (1:1000; ThermoFisher). F-actin was identified by Rhodamine-conjugated Phallodin (1:100; ThermoFisher) staining that was performed concurrently with secondary antibody incubations.

### Imaging and Statistical Analysis

Confocal imaging was done using an Olympus FluoView 1000 1000 (Figures 2AB, 3FIL, 6BCDEF, S3, S4, S10) within the University of Virginia Department of Cell Biology or a Zeiss LSM 700 (Figures 2C, 3, 4, 5, 6ABC, S1, S5, S6, S7, S8, S9, S12) in the University of Virginia Advanced Microscopy Facility (RRID:SCR_018736). Laser power and gain settings for each set of stained samples were based on the experimental group with the highest fluorescence intensity in each channel, and kept constant within the experiment. To compare between independently repeated experiments, we normalized within the experiment as indicated. Images were processed and quantified with Fiji/ImageJ (Schindelin et al., 2012).

Prism 8 software was used for Statistical Analysis. The specific tests that were used are listed in the figure descriptions.

### Dextran Assay

Larvae were inverted and cleaned in Schneider’s Insect Medium (Sigma-Aldrich) then transferred into a 1:8 dilution of 10 kD fluorescein conjugated dextran (Invitrogen) in Schneider’s Insect Medium (Sigma-Aldrich) and incubated rocking and covered at room temperature for 30 minutes. The tissues were washed briefly (approximately 1 minute) in Schneider’s Insect Medium (Sigma-Aldrich) to remove excess dextran, then fixed with 4% paraformaldehyde in Schnider’s Insect Media. Tissues were washed, stained, and imaged as previously described.

Fluorescent dextran infiltration was measured using Fiji/ImageJ (Schindelin et al., 2012), taking the mean intensity along a line in the imaginal disc lumen (identified by rhodamine phalloidin or Cora staining) and subtracting background from outside the disc area. Discs that appeared punctured were either not measured or categorized separately from intact discs. The fluorescence intensity varies with each experiment, so the data were normalized to the mean from controls that were incubated simultaneously.

### Quantification of septate junction component localization

Septate junction localization was quantified using Fiji/ImageJ (Schindelin et al., 2012). Two lines were drawn to collect fluorescence intensity of the junctions, the first across the apical-lateral surface of cells near the center of where the septate junctions were localized, and the second along the middle of the cells. Junctional intensity, or membrane intensity for the medial region, was considered as an average of the 7 pixels surrounding local maxima. In this way we hoped to average out misrepresentations in the data that arose from slices that cut through cells approximately parallel to the cell membranes and from slices that cut through tricellular junctions and had more protein from the third cell. None of the proteins we looked at are reported or appeared to have specific tricellular activity. We then took the ratio of the average junctional peak intensity to the average medial peak intensity (Fig. S7).

Peak identification was adjusted in three steps to reduce false identification of a membrane peak due to the noise within an image, especially with regards to Anti-Kune and Anti-Cora staining. First, to ensure the peak wasn’t a result of a slightly brighter random pixel, we removed peaks that were below the median fluorescence of the entire line. Second, to ensure the identified peak was localized at the membrane, we removed points of Anti-Kune and Anti-Cora staining that did not have a Nrx::GFP peak within the same 7-pixel range. We used Nrx::GFP instead of Actin (Rhodamine Phallodin) staining for this because Nrx::GFP has extremely low noise as it is a membrane bound GFP produced within the cell and does not need to be stained for. Nrx::GFP also has a very high association with the membrane even away from canonical apical-lateral staining, while this fluorescence is very dim, it is still detectable and highly correlated with the membrane. Finally, to ensure that we took a measurement at the membrane and not at a noisy region within the cell, if no Anti-Kune or Anti-Cora peak was identified within the 7-pixel range of the Nrx::GFP peak, a measurement was added at the same placement of the Nrx::GFP peak. Together these adjustments reduced the number of peak identifications in each image by approximately 1-10 junctions depending on the stain (most images had 40-60 junctions following adjustments).

## Acknowledgements

The authors would like to acknowledge Dr. David Bilder, Dr. Pierre Leopold, Dr. Maria Dominguez, Dr. Mikio Furuse, and Dr. Robert Ward for reagents and/or *Drosophila* stocks critical for the completion of these experiments. The authors would like to acknowledge the University of Virginia Advanced Microscopy Facility (RRID:SCR_018736) for training and access to the LSM700 confocal microscope.

## Data availability

There are no publicly available datasets associated with this work. All data underlying this work will be made available to researchers upon request.

---

**Figure S1.**
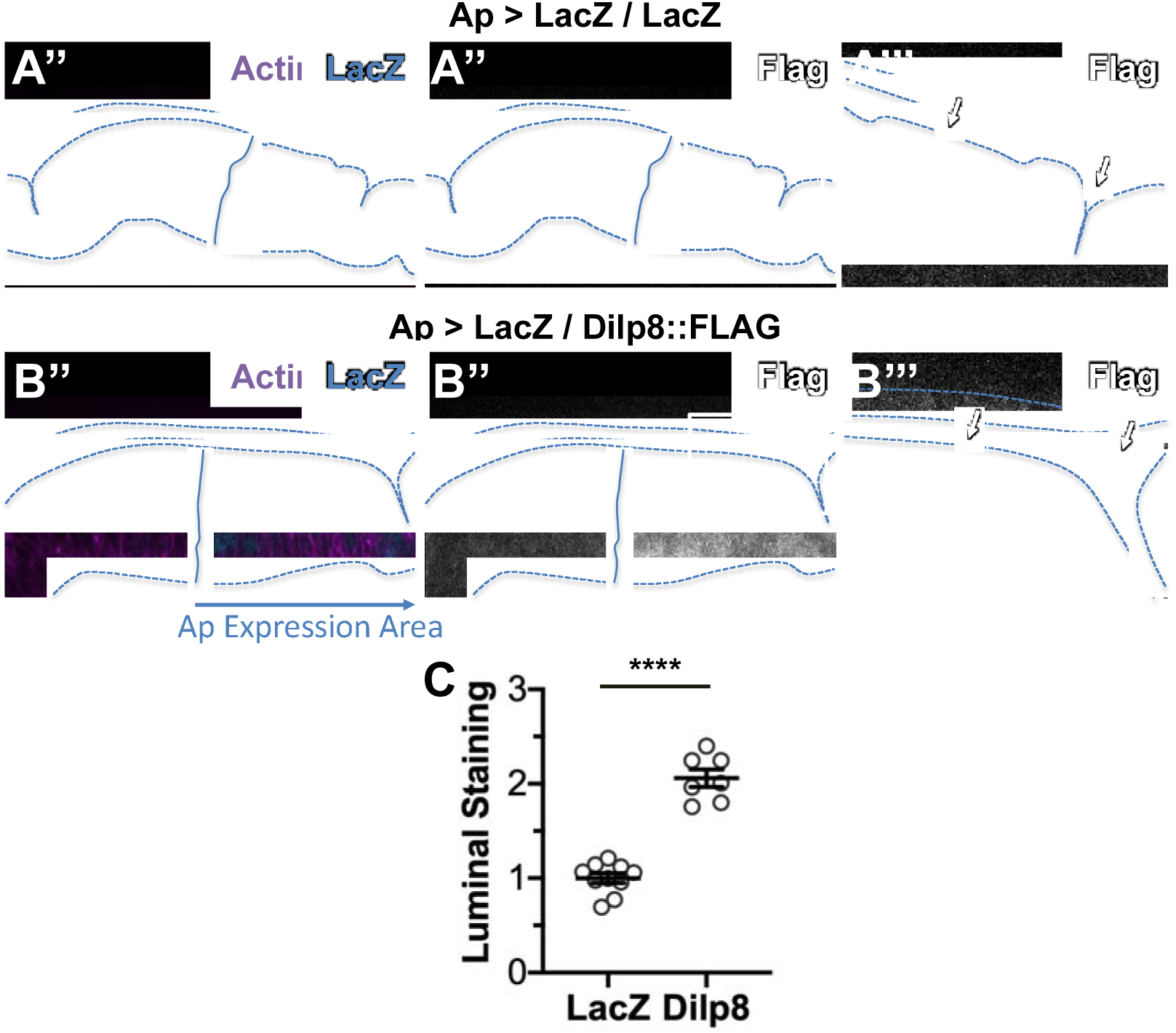
Dilp8 accumulates in the wing imaginal disc lumen. Ap-Gal4 was used to express (A) LacZ (wild type control) or (B) LacZ and Dilp8::FLAG in the dorsal region of wing imaginal discs. (A-B) Images are XZ cross-sections of wing imaginal discs in the pouch region of the disc. Dotted blue lines indicate disc area as defined by Actin (rhodamine phalloidin staining). Solid blue lines indicate the dorsal-ventral boundary, as defined by LacZ expression (β-Gal staining). Representative images are oriented dorsal on the right. (A) No FLAG is observed in *Ap > lacZ / lacZ* expressing discs either in (A’’) the expression region or (A’’’) the lumen. (B) FLAG is observed in *Ap > lacZ / dilp8::FLAG* expressing discs in the (B’’) expression region and (B’’’) the lumen. (C) Quantification of FLAG in the lumen, normalized to *Ap > LacZ* expression. Graph represent mean ± SEM, with individual points indicating values of single images. n = (LacZ) 10 and (Dilp8) 7 discs. **** p < 0.0001 by unpaired t-test.

**Figure S2.**
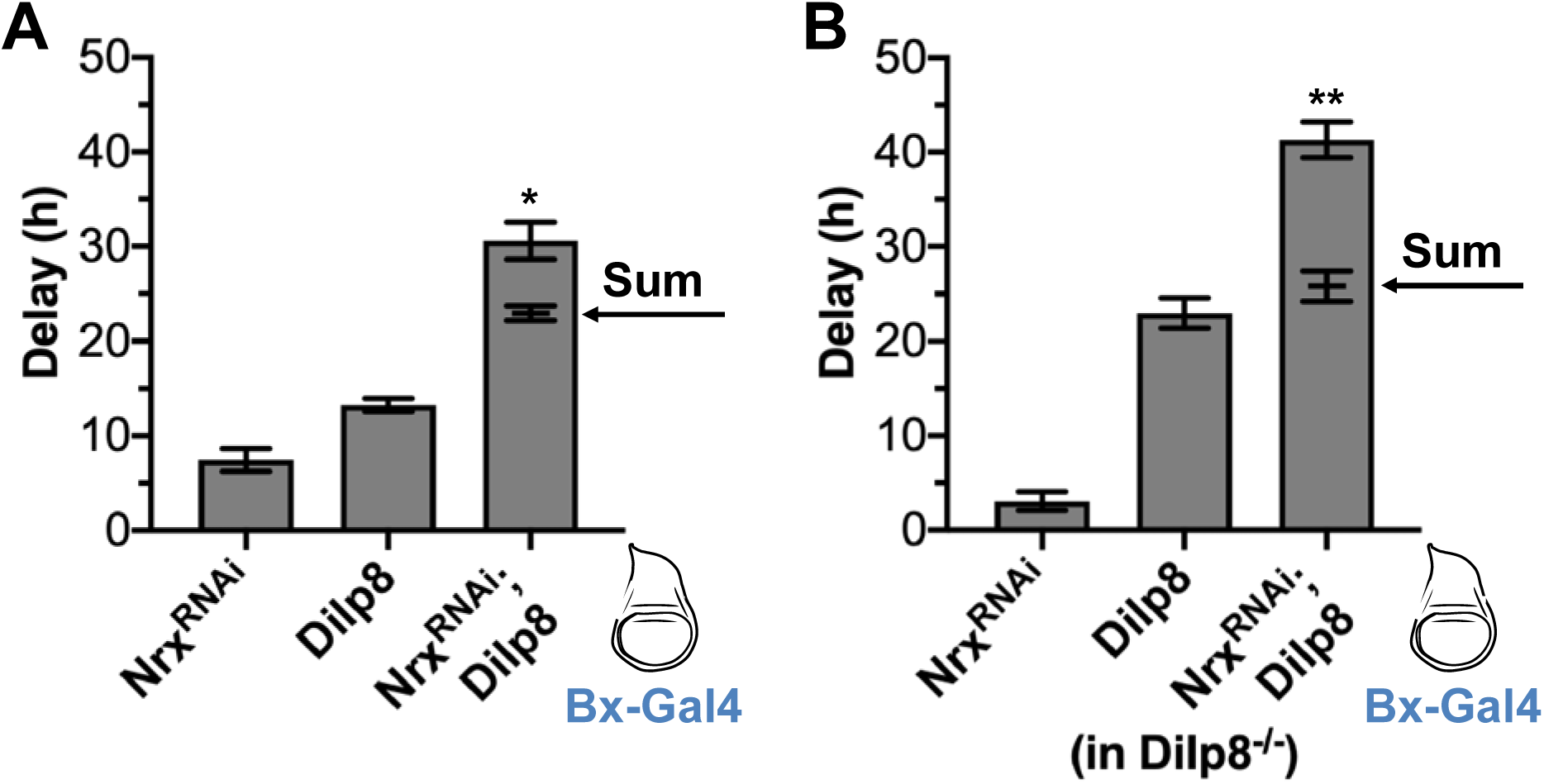
The epithelial barrier limits Dilp8 signaling. (A) Co-expression of *nrx*^*RNAi*^ and Dilp8 induces synergistic delay. Ectopic expression of nrx^RNAi^, Dilp8, and co-expression of *nrx*^*RNAi*^ and Dilp8 (*nrx*^*RNAi*^; Dilp8) induce developmental delay compared to LacZ controls when expressed in the wing imaginal disc under Bx-Gal4 (expression region in blue). The delay induced by co-expression of *nrx*^*RNAi*^ and Dilp8 (*nrx*^*RNAi*^; Dilp8) is significantly more than the sum of the delay induced by *nrx*^*RNAi*^ and Dilp8 expressed alone (sum indicated by arrow). (B) This trend holds true when endogenous Dilp8 is limited by expression in a Dilp8 hypomorphic background (Dilp8^MI00727^/Dilp8^MI00727^; Garelli et al., 2012). Data were collected from at least four independent experiments, bars represent mean ± SEM, * p < 0.05, ** p < 0.01 from one sample t-test comparing the additive value and observed delay.

**Figure S3.**
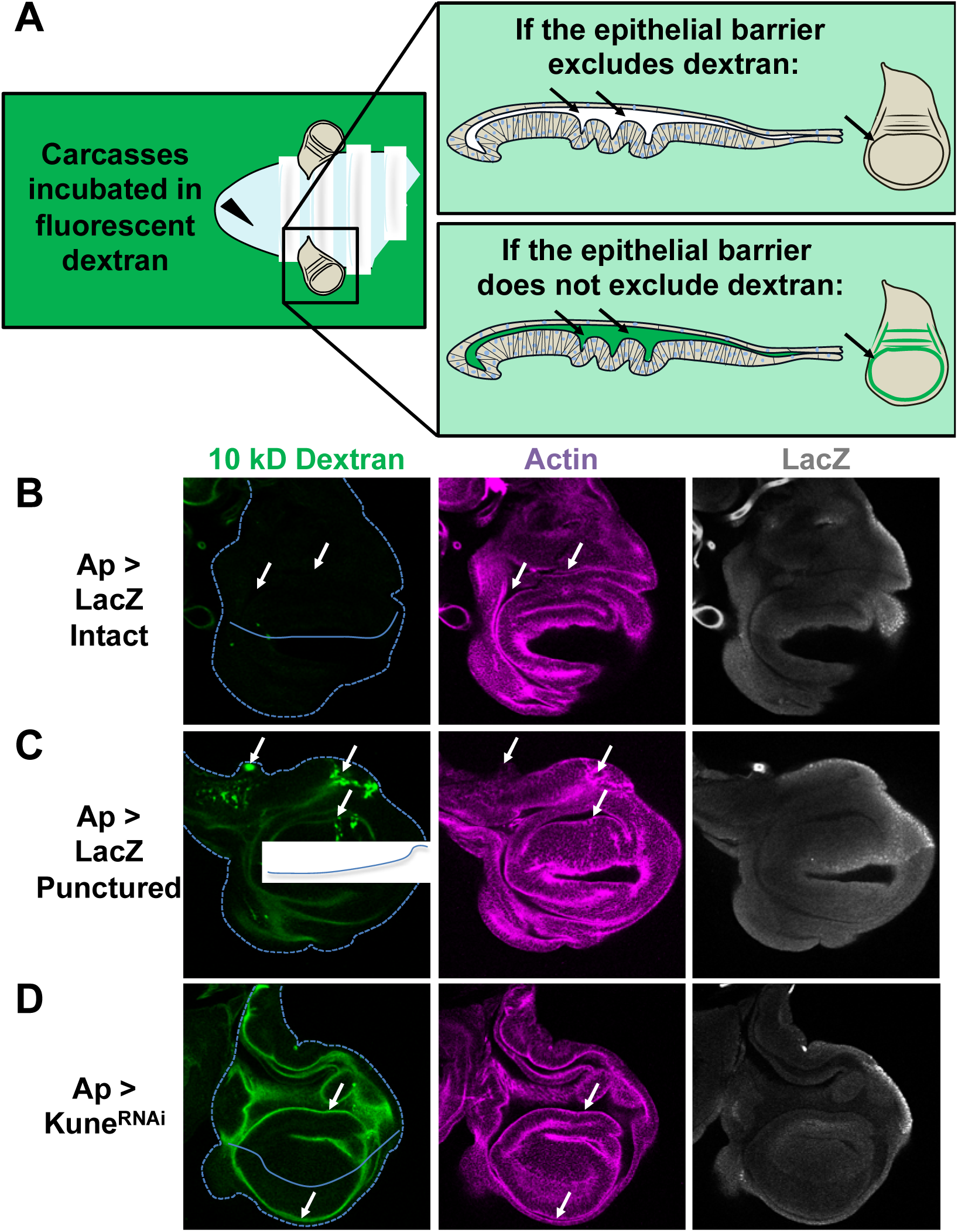
Explanation of the dextran assay for barrier function. (A) Carcasses are inverted and cleaned, then incubated in a fluorescence conjugated dextran for 30 minutes before fixation, the discs were then stained, mounted, and imaged (full description in methods). If the epithelial barrier excludes the dextran, no dextran should be observable in the lumen of the imaginal disc. If the epithelial barrier does not exclude the dextran or the tissue integrity is disrupted, dextran should be observable in the lumen. (B-D) Representative images after 10 kD fluorescein-conjugated dextran incubation (from the experiment quantified in Figure 2A). Ap-Gal4 was used to express LacZ (Ap > LacZ) or Kune^RNAi^ (Ap > Kune^RNAi^). Dextran is observed in discs punctured LacZ expressing discs (C) and Kune^RNAi^ expressing discs (D), but not in intact LacZ expressing discs (A). Disc area is indicated by the dashed line, as defined by Actin staining (rhodamine phalloidin). Area of expression is dorsal (oriented up) of the solid line, as defined by LacZ staining (b-Gal; note that LacZ expressing discs express two copies of LacZ while Kune^RNAi^ expressing discs have one copy so β-Gal staining is not comparable between images). Arrows indicate: (A) areas of the lumen with no distinguishable dextran fluorescence; (B) areas where damage during dissection (puncturing) has disrupted epithelial barrier integrity, allowing dextran to enter the cells and the imaginal disc lumen; and (C) luminal dextran is observed in both the dorsal lumen (Kune^RNAi^ expressing area, oriented up) and also in the ventral lumen (non-expressing area, oriented down), indicating that the lumen is contiguous. Images are single slices.

**Figure S4.**
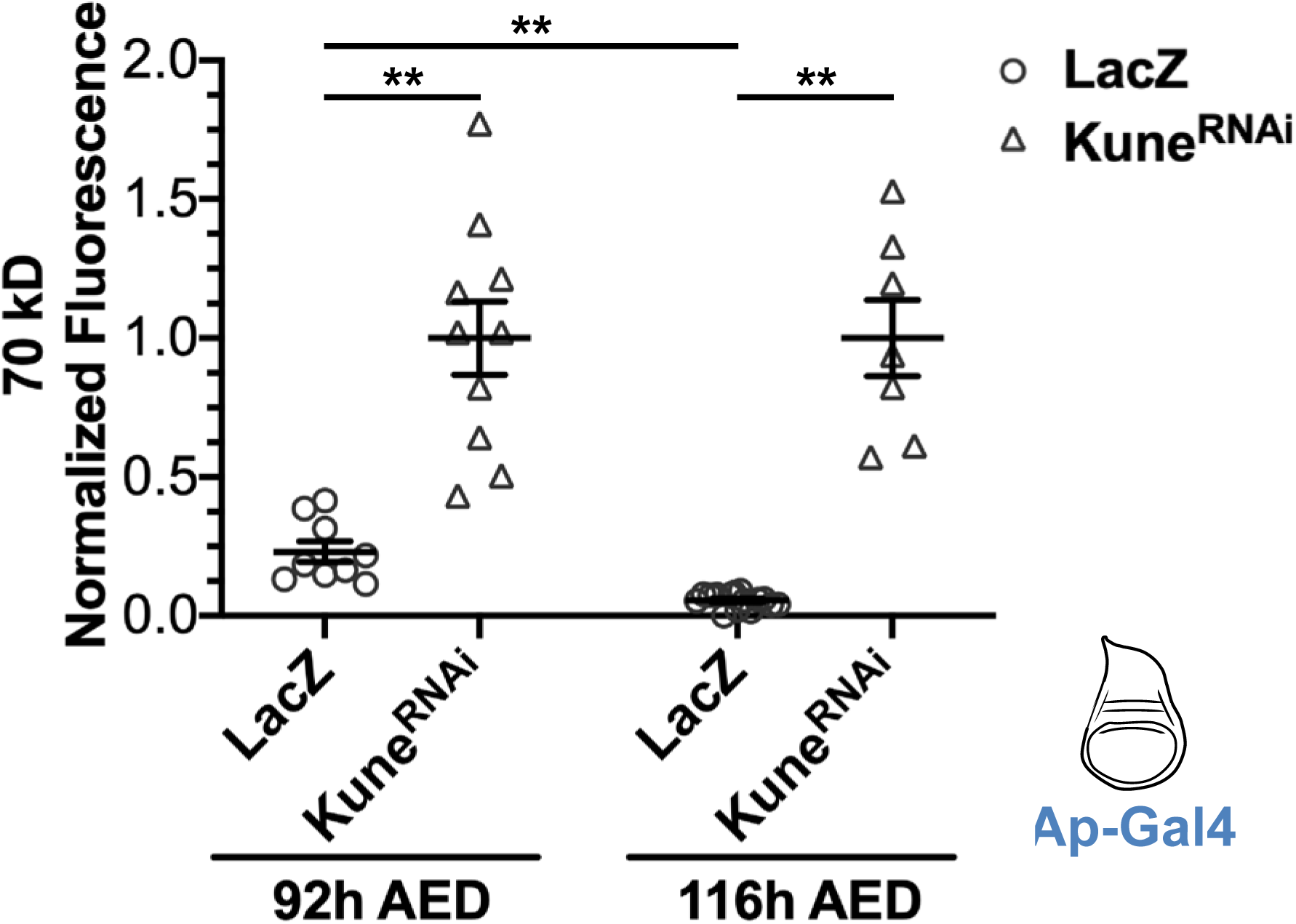
The epithelial barrier of wing imaginal discs grow more exclusionary to 70 kD fluorescent dextran during the third instar. The function of the epithelial barrier to exclude 70 kD Texas Red conjugated dextran was measured, as previously described, at 92h and 116h AED in wing imaginal discs expressing LacZ or Kune^RNAi^ by Ap-Gal4 (expression area diagramed in blue). Data are normalized to the mean luminal intensity of the Kune^RNAi^ expressing discs. Graph represents mean ± SEM, with individual points indicating values of single images. Left to right, n = 9, 10, 15, 7. ** p < 0.01 as calculated by Brown-Forsythe and Welch ANOVA with Dunnett’s T3 test for multiple comparisons.

**Figure S5.**
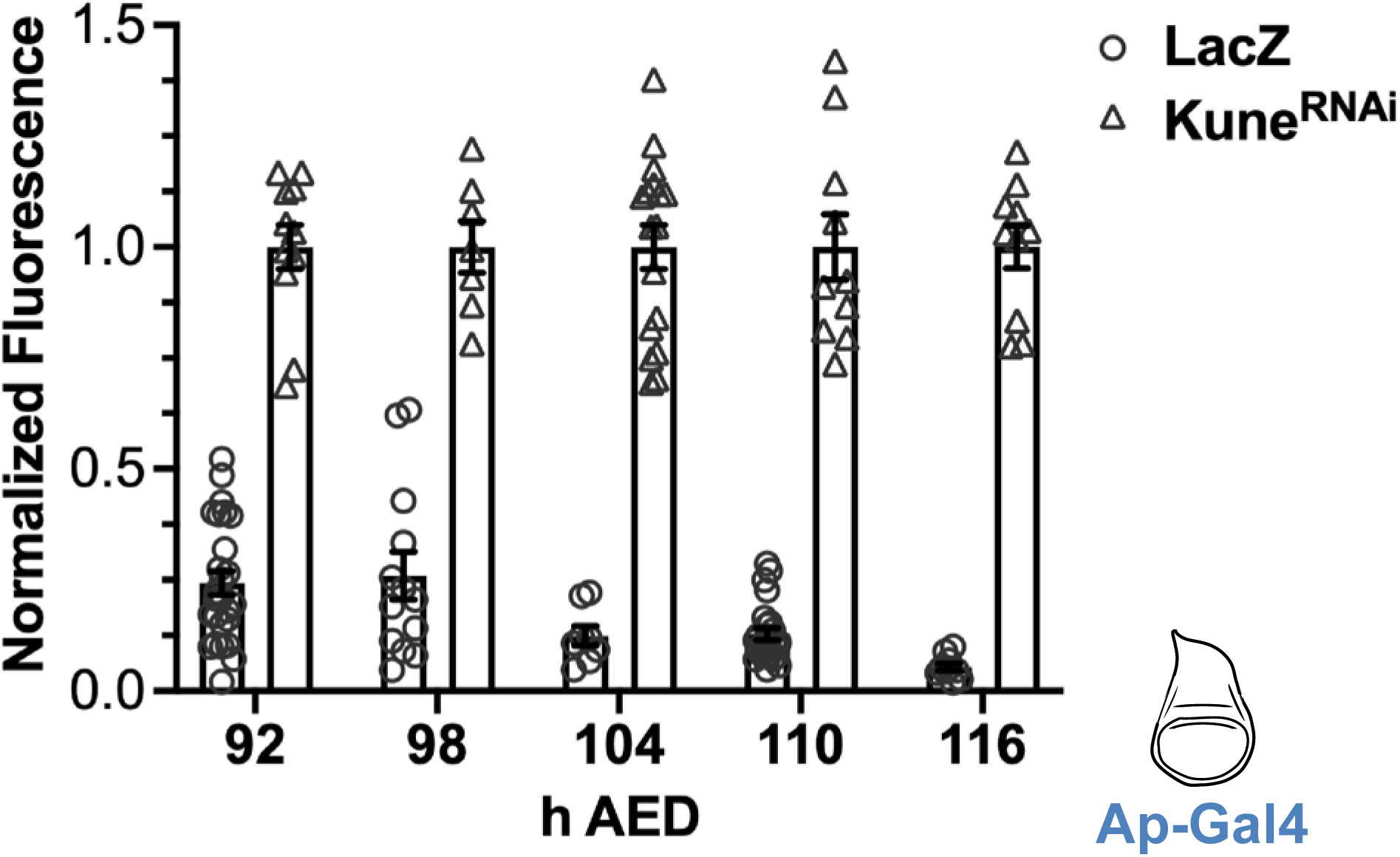
The epithelial barrier matures between 92h and 116h AED, becoming more restrictive to 10 kD fluorescein conjugated dextran. These are the complete data from Figure 2C, including data from Kune^RNAi^ expressing discs: the barrier function of wing imaginal discs expressing LacZ or Kune^RNAi^ by Ap-Gal4 was measured every 6 hours between 92h and 116h AED. Data indicate luminal intensity of intact LacZ expressing discs normalized to the mean luminal intensity of the Kune^RNAi^ expressing discs from the same timepoint. Graphs represent mean ± SEM, with individual points indicating values of single images. Significance between LacZ expressing discs at each timepoint is indicated in Figure 2C. Left to right, n = (92h AED) 26, 11, (98h AED) 13, 7, (104h AED) 12, 17, (110h AED) 25, 10, (116h AED) 10, 11.

**Figure S6.**
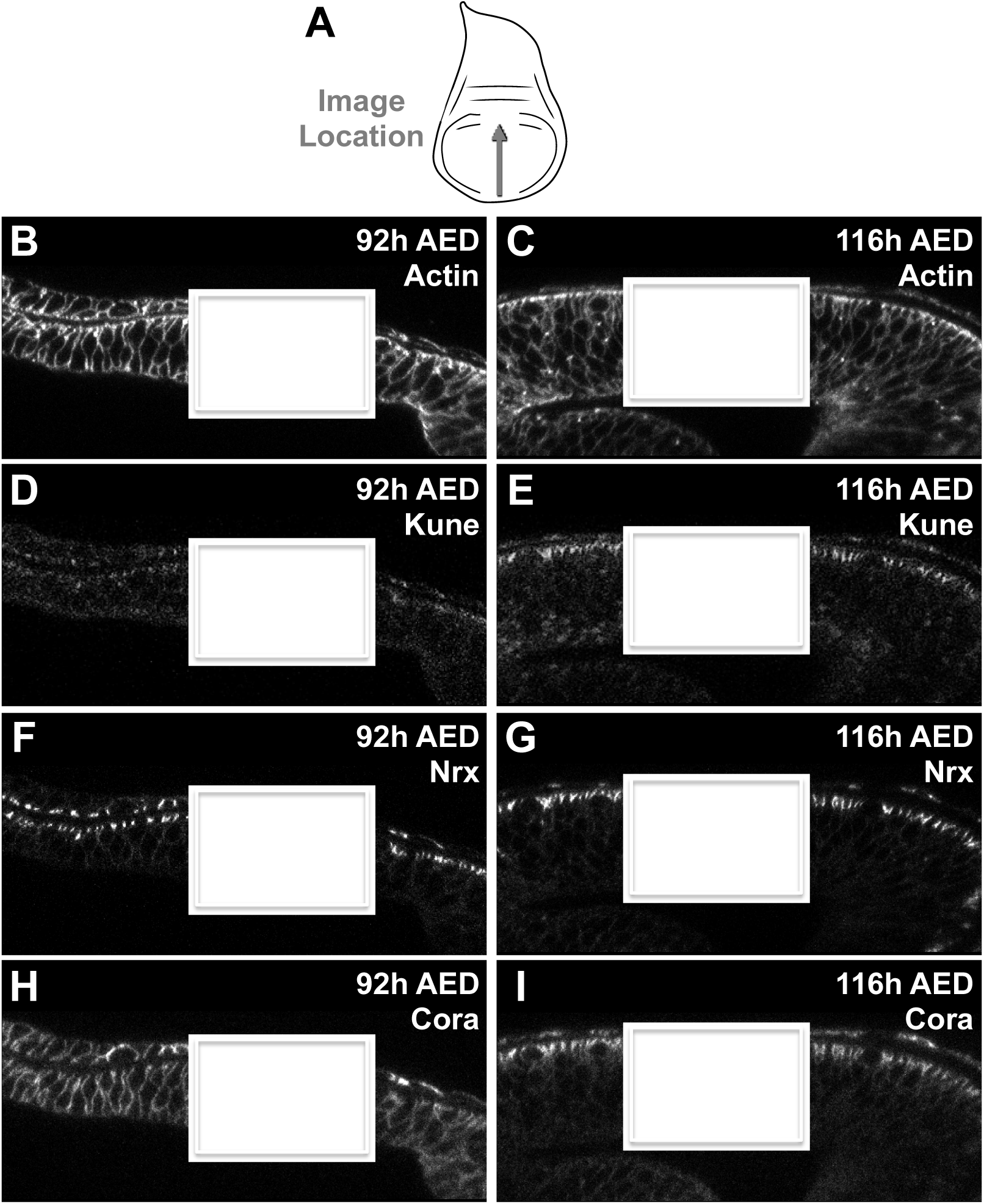
Complete images from Figure 3. (A) Approximate area and orientation of XZ image locations in the wing imaginal discs. (B-I) Representative XZ images at 92h and 116h AED, the region zoomed into in Figure 3 is indicated (white box). Images are: (B-C) Actin (rhodamine phalloidin; corresponds to Figure 3 B-C), (D-E) Kune (Anti-Kune; corresponds to Figure 3 E-F), (F-G) Nrx (Nrx-GFP; corresponds to Figure 3 G-H); and (H-I) Cora (Anti-Cora; corresponds to Figure 3 J-K).

**Figure S7.**
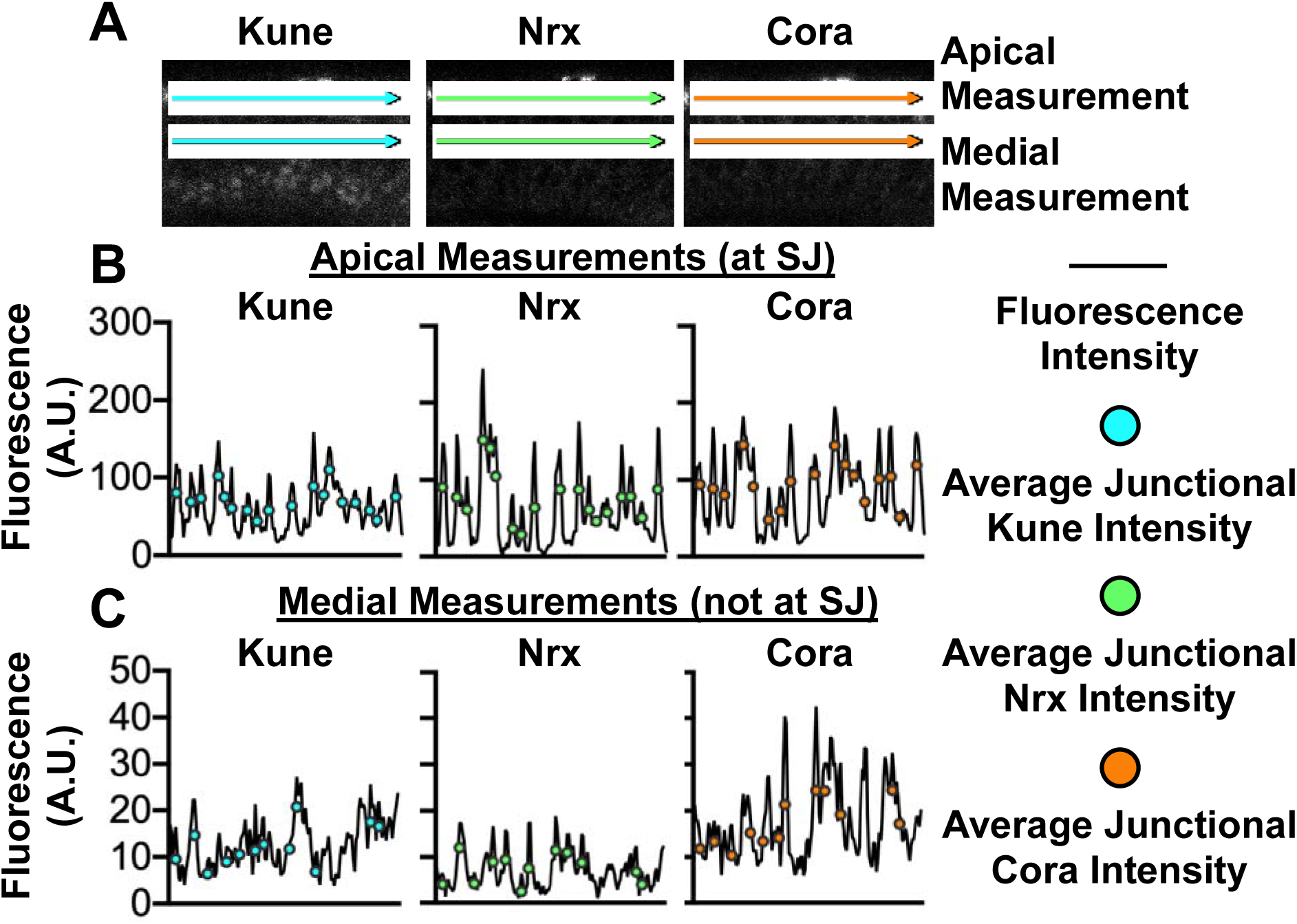
Method for junction quantification. (A) Lines were drawn apically bisecting the region of brightest septate junction (SJ) staining, and medially. (B-C) Fluorescence intensity across each line were measured with plot profile. Areas of the membrane (septate junction if apical) were identified as local maxima (peak) within a 7-pixel range, the 3 prior and following the pixel in questions. We adjusted these data in three steps, further described in Materials and Methods, to reduce false identification of a peak being localized at a membrane due to imaging or staining issues (eg. non-specific staining, image noise), especially with regards to Anti-Kune and Anti-Cora staining. The average junctional intensity was taken as the mean of the 7-pixel range.

**Figure S8.**
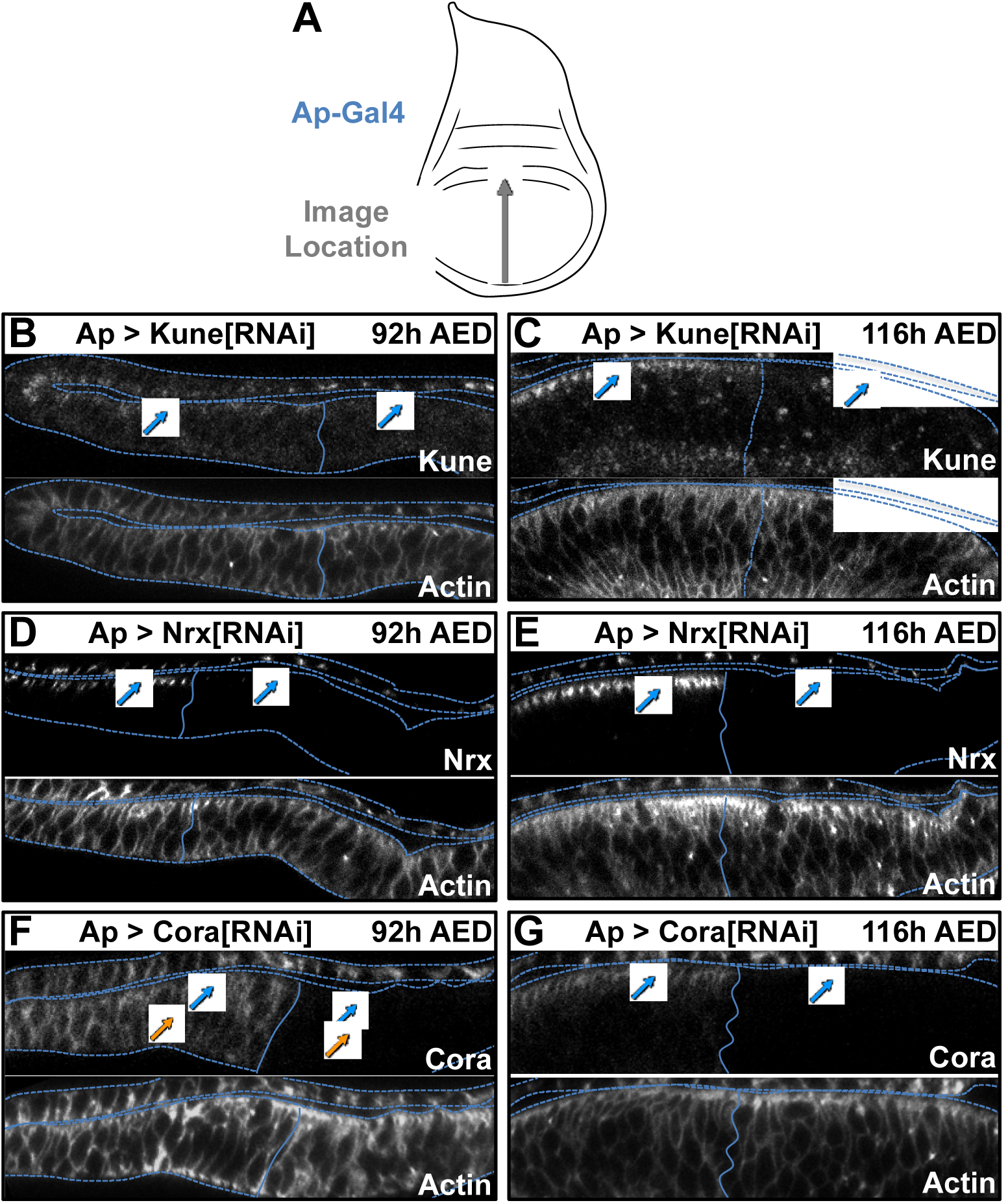
RNAi inhibition of septate junction components. (A) Images were taken spanning the dorsal-ventral boundary in the pouch region of wing imaginal discs (approximate area indicated by grey arrow), which includes tissue outside and inside the Ap-Gal4 expression region (blue). (B-G) Localization of septate junction components following RNAi expression and actin localization (defined by rhodamine phalloidin). Images are from the same discs as Figure 5. Dotted line represents tissue outline defined by actin staining. Solid line represents dorsal-ventral boundary, expression area (dorsal region) is on the right. Blue arrows indicate apical-lateral localization, orange arrows indicate medial-lateral localization. (B-C) Localization of Kune and Actin in *Ap > kune*^*RNAi*^ expressing discs at (B) 92h and (C) 116h AED. Kune is depleted in the *kune*^*RNAi*^ expression region at both times. Images are of the same discs as Figure 4A and 4B. (D-E) Localization of Nrx and Actin in *Ap > nrx*^*RNAi*^ expressing discs at (D) 92h and (E) 116h AED. Nrx is depleted in the *nrx*^*RNAi*^ expression region at both times. Images are of the same discs as Figure 4C and 4D. (F-G) Localization of Cora and Actin in *Ap > cora*^*RNAi*^ expressing discs at (F) 92h and (G) 116h AED. Kune is depleted in the *cora*^*RNAi*^ expression region at both times. Images are of the same discs as Figure 4E and 4F.

**Figure S9.**
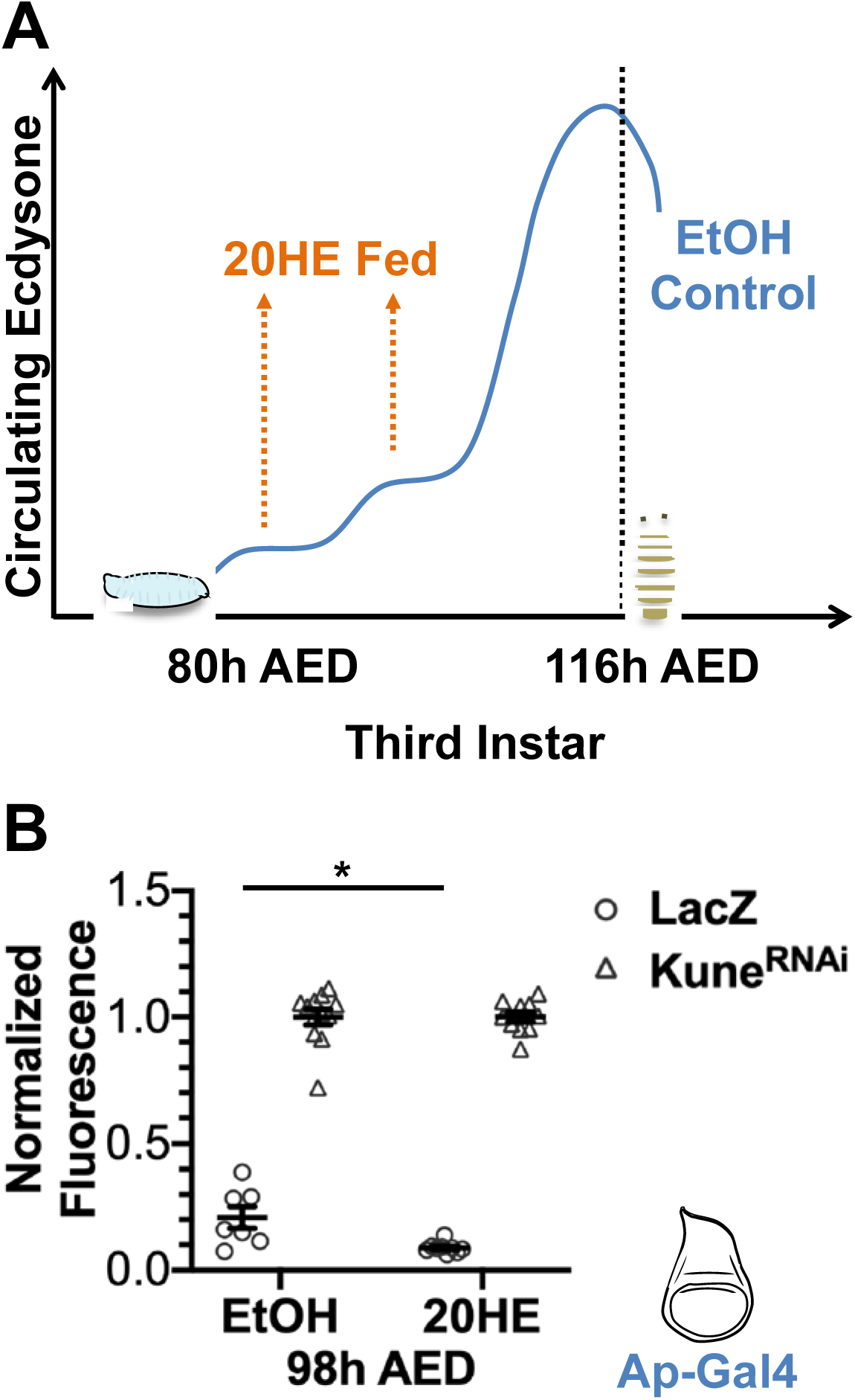
Ecdysone feeding induces barrier maturation early. (A) At 80h AED, larvae were switched to food containing 0.6 mg/mL 20-hydroxyecdysone (20HE) or ethanol control (EtOH). Feeding 0.6 mg/mL 20HE at 80h AED does not influence pupariation time, but does limit regeneration (Colombani et al., 2005; Jaszczak et al., 2015). (B) Wing imaginal disc barrier function at 98h AED of larvae fed the EtOH or 20HE food. Data are from larvae expressing *Ap > lacZ* (data in Figure 6A) or *Ap > kune*^*RNAi*^. Expression area indicated in blue. Graph represents mean ± SEM, with individual points indicating values of single images. Left to right, n = 7, 12, 9, 10. * p < 0.05 as calculated by unpaired t-test with Welch’s correction.

**Figure S10.**
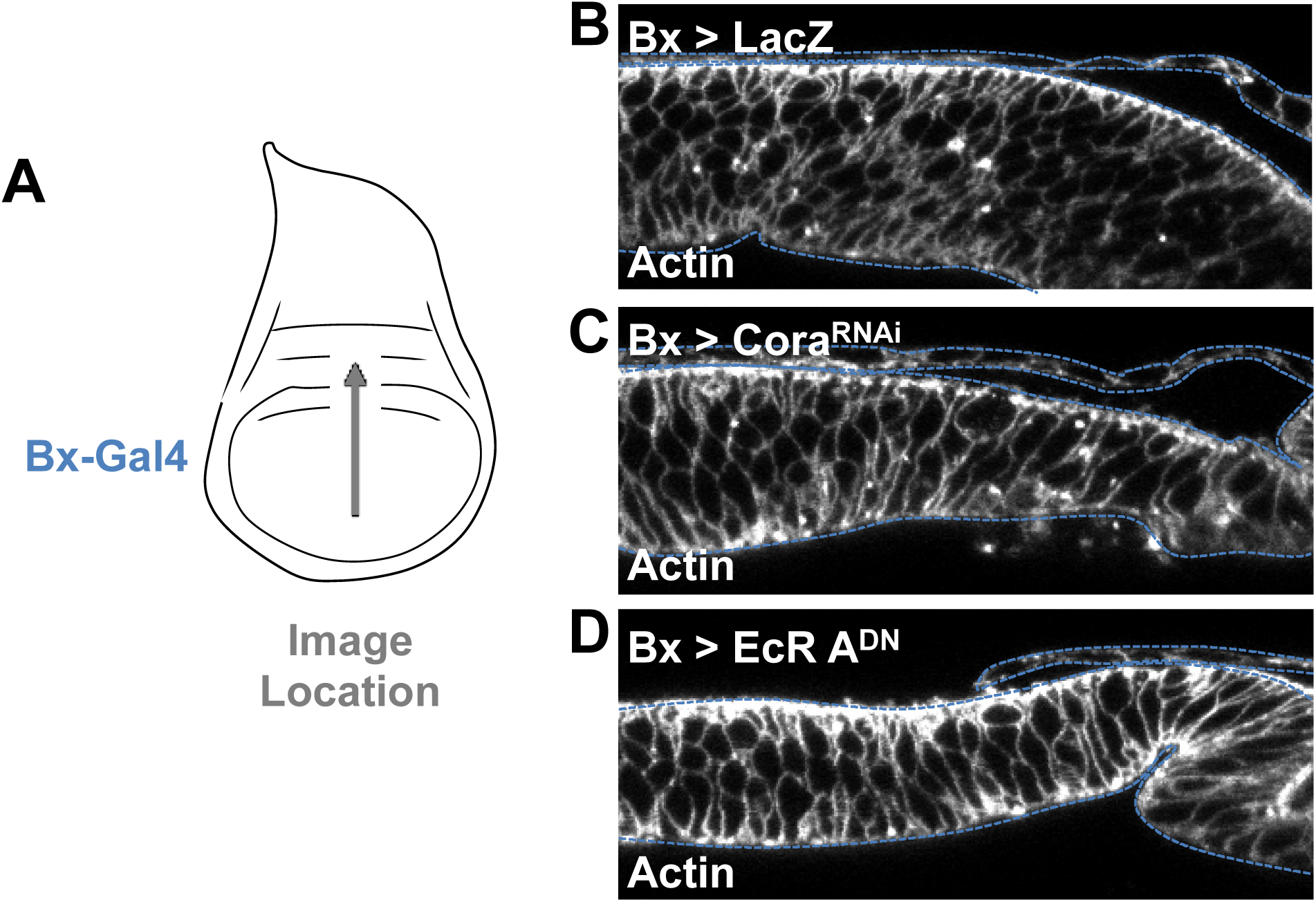
Actin stain from discs in Figure 6. (A) Bx-Gal4 expression area (blue) and approximate image location (grey arrow) for B-D. (B-D) Localization of Actin (rhodamine phalloidin) in (B) *Bx > lacZ* (wild type control), (C) *Bx > kune*^*RNAi*^, and (D) *Bx > EcR A*^*DN*^ at 116h AED. Dotted lines indicate tissue outline. Tissues are oriented with dorsal on the right. Images are from the same discs as Figure 6D-F.

**Figure S11.**
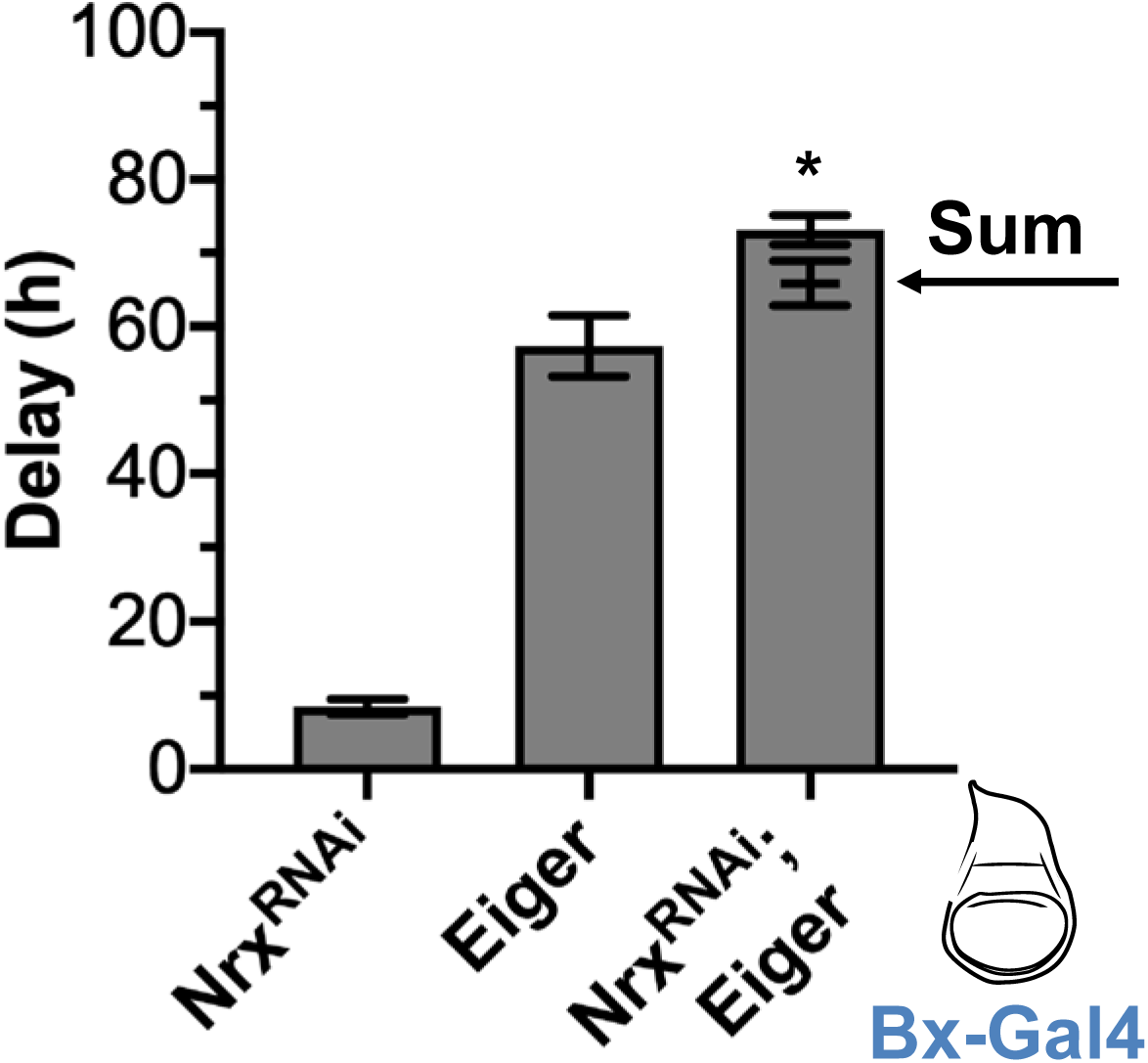
Nrx limits Eiger-induced delay. Co-expression of *nrx*^*RNAi*^ and Eiger produces synergistic delay. Ectopic expression of *nrx*^*RNAi*^, Eiger, and co-expression of *nrx*^*RNAi*^ and Eiger (*nrx*^*RNAi*^; Eiger) induce developmental delay compared to LacZ controls when expressed in the wing imaginal disc under Bx-Gal4 (expression region in blue). The delay induced by co-expression of *nrx*^*RNAi*^ and Eiger (*nrx*^*RNAi*^; Eiger) is significantly more than the sum of the delay induced by *nrx*^*RNAi*^ and Eiger expressed alone (sum indicated by arrow. Data were collected from at least four independent experiments, bars represent mean ± SEM, * p < 0.01 from one sample t-test comparing the additive value and observed delay.

**Figure S12.**
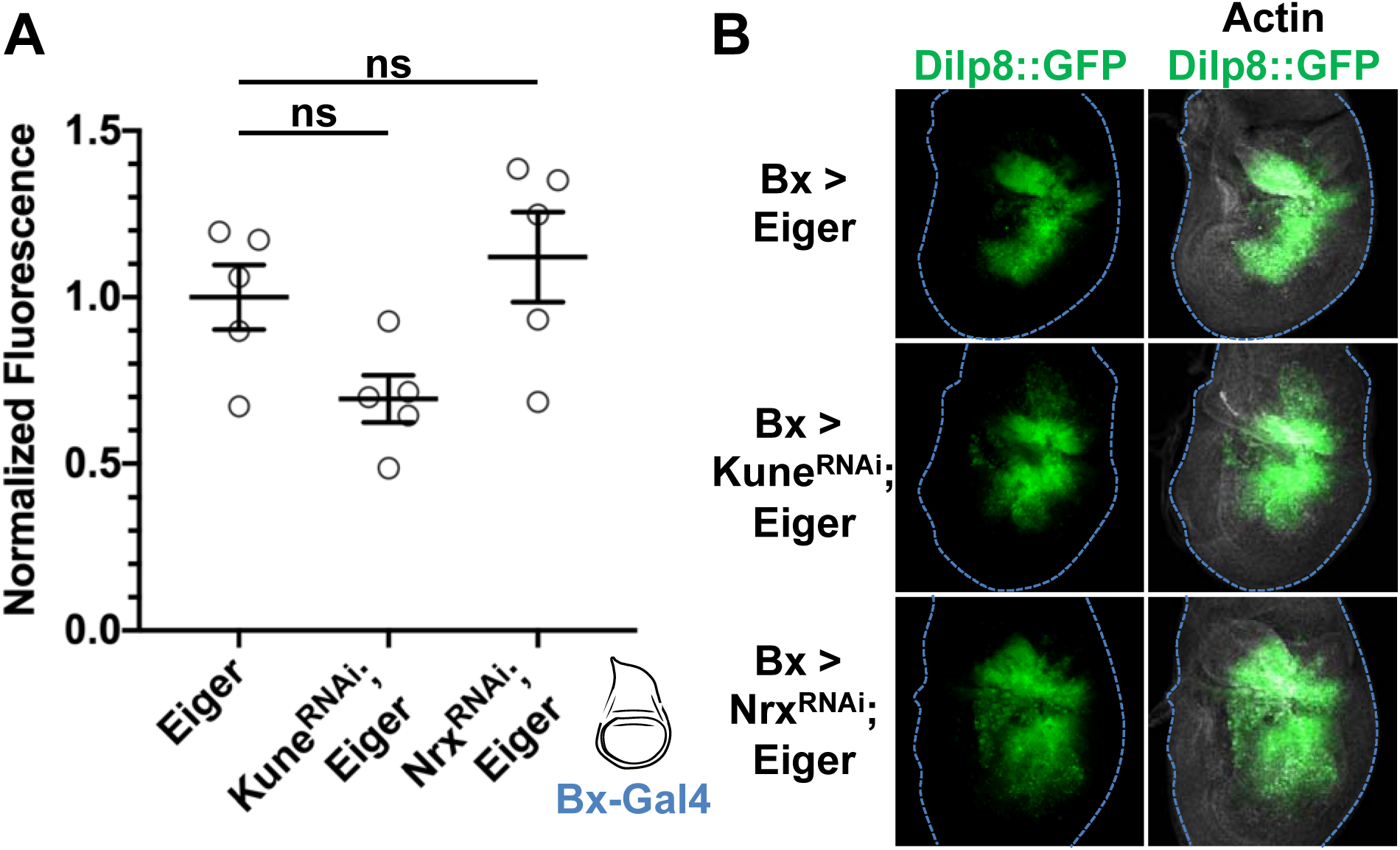
Coexpression of Eiger and RNAi against septate junction components does not significantly alter measured Dilp8 expression. Bx-Gal4 (expression area in blue) was used to express Eiger alone, with *kune*^*RNAi*^, or with *nrx*^*RNAi*^ expressing a transcriptional reporter for Dilp8 (Dilp8^MI00727^/+; Garelli et al., 2012). (A) Sum GFP intensity was measured and the data were normalized to Eiger alone. (B) Representative images. Discs were collected at 104 hAED. Images represent sum-projected stacks of 5 images. Actin was stained for with rhodamine phalloidin. (A) Graph represents mean ± SEM, with individual points indicating values of single images, n = 5 images in each condition; ns indicates p > 0.05 by one-way ANOVA with Tukey test for multiple comparisons.

## References

Baumgartner, S., Littleton, J. T., Broadie, K., Bhat, M. a., Harbecke, R., Lengyel, J. a., Chiquet-Ehrismann, R., Prokop, A. and Bellen, H. J. (1996). A Drosophila neurexin is required for septate junction and blood-nerve barrier formation and function. Cell 87, 1059–1068.

Buszczak, M., Paterno, S., Lighthouse, D., Bachman, J., Planck, J., Owen, S., Skora, A. D., Nystul, T. G., Ohlstein, B., Allen, A., et al. (2007). The carnegie protein trap library: A versatile tool for drosophila developmental studies. Genetics 175, 1505–1531.

Cohen, B., McGuffin, M. E., Pfeifle, C., Segal, D. and Cohen, S. M. (1992). apterous, a gene required for imaginal disc development in Drosophila encodes a member of the LIM family of developmental regulatory proteins. Genes and Development 6, 715–729.

Colombani, J., Bianchini, L., Layalle, S., Pondeville, E., Dauphin-Villemant, C., Antoniewski, C., Carré, C., Noselli, S. and Léopold, P. (2005). Antagonistic actions of ecdysone and insulins determine final size in Drosophila. Science 310, 667–670.

Colombani, J., Andersen, D. S. and Leopold, P. (2012). Secreted Peptide Dilp8 Coordinates Drosophila Tissue Growth with Developmental Timing. Science 336, 582–585.

Colombani, J., Andersen, D. S., Boulan, L., Boone, E., Romero, N., Virolle, V., Texada, M. and Léopold, P. (2015). Drosophila Lgr3 Couples Organ Growth with Maturation and Ensures Developmental Stability. Current biology□: CB 25, 2723–9.

Fehon, R. G., Dawson, I. a and Artavanis-Tsakonas, S. (1994). A Drosophila homologue of membrane-skeleton protein 4.1 is associated with septate junctions and is encoded by the coracle gene. Development (Cambridge, England) 120, 545–557.

Fox, D. T., Cohen, E. and Smith-Bolton, R. (2020). Model systems for regeneration: Drosophila. Development 147,.

Furuse, M. and Tsukita, S. (2006). Claudins in occluding junctions of humans and flies. Trends in cell biology 16, 181–8.

Garelli, a., Gontijo, a. M., Miguela, V., Caparros, E. and Dominguez, M. (2012). Imaginal Discs Secrete Insulin-Like Peptide 8 to Mediate Plasticity of Growth and Maturation. Science 336, 579–582.

Garelli, A., Heredia, F., Casimiro, A. P., Macedo, A., Nunes, C., Garcez, M., Mantas Dias, A. R., Volonte, Y. A., Uhlmann, T., Caparros, E., et al. (2015). ARTICLE Dilp8 requires the neuronal relaxin receptor Lgr3 to couple growth to developmental timing.

Genova, J. L. and Fehon, R. G. (2003). Neuroglian, Gliotactin, and the Na+/k+ ATPase are essential for septate junction function in Drosophila. Journal of Cell Biology 161, 979–989.

Hackney, J. F. and Cherbas, P. (2014). Injury response checkpoint and developmental timing in insects. Fly 8, 226–231.

Hackney, J. F., Zolali-Meybodi, O. and Cherbas, P. (2012). Tissue damage disrupts developmental progression and ecdysteroid biosynthesis in Drosophila. PloS one 7, e49105.

Hall, S. and Ward, R. E. (2016). Septate Junction Proteins Play Essential Roles in Morphogenesis Throughout Embryonic Development in Drosophila. G3 (Bethesda, Md.) 6, 2375–84.

Halme, A., Cheng, M. and Hariharan, I. K. (2010). Retinoids Regulate a Developmental Checkpoint for Tissue Regeneration in Drosophila. Current Biology 20, 458–463.

Hamaratoglu, F., Affolter, M. and Pyrowolakis, G. (2014). Dpp/BMP signaling in flies: From molecules to biology. Seminars in Cell & Developmental Biology 32, 128–136.

Igaki, T., Kanda, H., Yamamoto-Goto, Y., Kanuka, H., Kuranaga, E., Aigaki, T. and Miura, M. (2002). Eiger, a TNF superfamily ligand that triggers the Drosophila JNK pathway. EMBO Journal 21, 3009–3018.

Izumi, Y. and Furuse, M. (2014). Molecular organization and function of invertebrate occluding junctions. Seminars in Cell & Developmental Biology 36, 186–193.

Jaszczak, J. S., Wolpe, J. B., Dao, A. Q. and Halme, A. (2015). Nitric oxide synthase regulates growth coordination during Drosophila melanogaster imaginal disc regeneration. Genetics 200, 1219–1228.

Jaszczak, J. S., Wolpe, J. B., Bhandari, R., Jaszczak, R. G. and Halme, A. (2016). Growth Coordination During Drosophila melanogaster Imaginal Disc Regeneration Is Mediated by Signaling Through the Relaxin Receptor Lgr3 in the Prothoracic Gland. Genetics 204, 703–709.

Kauppila, S., Maaty, W. S. A., Chen, P., Tomar, R. S., Eby, M. T., Chapo, J., Chew, S., Rathore, N., Zachariah, S., Sinha, S. K., et al. (2003). Eiger and its receptor, Wengen, comprise a TNF-like system in Drosophila. Oncogene 22, 4860–4867.

Kozlova, T. and Thummel, C. S. (2000). Steroid Regulation of Postembryonic Development and Reproduction in Drosophila. Trends in Endocrinology & Metabolism 11, 276–280.

Kozlova, T. and Thummel, C. S. (2003). Essential Roles for Ecdysone Signaling During Drosophila Mid-Embryonic Development. Science 301, 1911–1914.

Lamb, R. S., Ward, R. E., Schweizer, L. and Fehon, R. G. (1998). Drosophila coracle, a member of the protein 4.1 superfamily, has essential structural functions in the septate junctions and developmental functions in embryonic and adult epithelial cells. Molecular biology of the cell 9, 3505–3519.

Milán, M., Diaz-Benjumea, F. J. and Cohen, S. M. (1998). Beadex encodes an LMO protein that regulates Apterous LIM-homeodomain activity in Drosophila wing development: A model for LMO oncogene function. Genes and Development 12, 2912–2920.

Moreno, E., Yan, M. and Basler, K. (2002). Evolution of TNF signaling mechanisms: JNK-dependent apoptosis triggered by Eiger, the Drosophila homolog of the TNF superfamily. Current Biology 12, 1263–1268.

Morin, X., Daneman, R., Zavortink, M. and Chia, W. (2001). A protein trap strategy to detect GFP-tagged proteins expressed from their endogenous loci in Drosophila. Proceedings of the National Academy of Sciences of the United States of America 98, 15050–15055.

Nelson, K. S., Furuse, M. and Beitel, G. J. (2010). The Drosophila Claudin Kune-kune is required for septate junction organization and tracheal tube size control. Genetics 185, 831–9.

Oshima, K. and Fehon, R. G. (2011). Analysis of protein dynamics within the septate junction reveals a highly stable core protein complex that does not include the basolateral polarity protein Discs large. Journal of Cell Science 124, 2861–2871.

Pastor-Pareja, J. C., Grawe, F., Martín-Blanco, E. and García-Bellido, A. (2004). Invasive cell behavior during Drosophila imaginal disc eversion is mediated by the JNK signaling cascade. Developmental Cell 7, 387–399.

Paul, S. M., Ternet, M., Salvaterra, P. M. and Beitel, G. J. (2003). The Na+/K+ ATPase is required for septate junction function and epithelial tube-size control in the Drosophila tracheal system. Development (Cambridge, England) 130, 4963–4974.

Schindelin, J., Arganda-Carreras, I., Frise, E., Kaynig, V., Longair, M., Pietzsch, T., Preibisch, S., Rueden, C., Saalfeld, S., Schmid, B., et al. (2012). Fiji: an open-source platform for biological-image analysis. Nature Methods 9, 676–682.

Setiawan, L., Pan, X., Woods, A. L., O’connor, M. B. and Hariharan, I. K. (2018). The BMP2/4 ortholog Dpp can function as an inter-organ signal that regulates developmental timing.

Tan, K. L., Vlisidou, I. and Wood, W. (2014). Ecdysone Mediates the Development of Immunity in the Drosophila Embryo. Current Biology 24, 1145–1152.

Tepass, U., Tanentzapf, G., Ward, R. and Fehon, R. (2001). EPITHELIAL CELL POLARITY AND CELL JUNCTIONS IN DROSOPHILA.

Vallejo, D. M., Juarez-Carreño, S., Bolivar, J., Morante, J. and Dominguez, M. (2015). A brain circuit that synchronizes growth and maturation revealed through Dilp8 binding to Lgr3. Science (New York, N.Y.) 1–16.

Ward IV, R. E., Schweizer, L., Lamb, R. S. and Fehon, R. G. (2001). The protein 4.1, ezrin, radixin, moesin (FERM) domain of drosophila coracle, a cytoplasmic component of the septate junction, provides functions essential for embryonic development and imaginal cell proliferation. Genetics 159, 219–228.

