## Supplemental information and figures for "Ecdysone regulates the *Drosophila* imaginal disc epithelial barrier, determining the duration of regeneration checkpoint delay"

**Supplementary Materials and Methods**

**Genotypes**

Figure 1A

Bx-Gal4 / + ; UAS-Dcr2 / + ; UAS-LacZ.NZ / +

Bx-Gal4 / + ; UAS-Dcr2 / UAS-Kune^RNAi^

Bx-Gal4 / + ; UAS-Dcr2 / + ; UAS-Dilp8::3xFLAG / +

Bx-Gal4 / + ; UAS-Dcr2 / UAS-Kune^RNAi^ ; UAS-Dilp8::3xFLAG / +

Figure 1B

Bx-Gal4 / + ; UAS-Dcr2 / + ; Dilp8^MI00727^ / Dilp8^MI00727^

Bx-Gal4 / + ; UAS-Dcr2 / UAS-Kune^RNAi^ ; Dilp8^MI00727^ / Dilp8^MI00727^

Bx-Gal4 / + ; UAS-Dcr2 / + ; UAS-Dilp8::3xFLAG , Dilp8^MI00727^ / Dilp8^MI00727^

Bx-Gal4 / + ; UAS-Dcr2 / UAS-Kune^RNAi^ ; UAS-Dilp8::3xFLAG , Dilp8^MI00727^ / Dilp8^MI00727^

Figure 2

Ap-Gal4 / + ; UAS-Dcr2 , UAS-LacZ.NZ / UAS-LacZ.NZ

Ap-Gal4 / UAS-Kune^RNAi^ ; UAS-Dcr2 , UAS-LacZ.NZ / +

Figure 3

NrxIV-GFP / NrxIV-GFP

Figure 4

Ap-Gal4 / + ; UAS-Dcr2 , UAS-LacZ.NZ / UAS-LacZ.NZ

Ap-Gal4 / UAS-Kune^RNAi^ ; UAS-Dcr2 , UAS-LacZ.NZ / +

Ap-Gal4 / UAS-Cora^RNAi^ ; UAS-Dcr2 , UAS-LacZ.NZ / +

Figure 5A,B

Ap-Gal4 / UAS-Kune^RNAi^ ; UAS-Dcr2 , UAS-LacZ.NZ / NrxIV-GFP

Figure 5C,D

Ap-Gal4 / UAS-Nrx^RNAi^ ; UAS-Dcr2 , UAS-LacZ.NZ / NrxIV-GFP

Figure 5E,F

Ap-Gal4 / UAS-Cora^RNAi^ ; UAS-Dcr2 , UAS-LacZ.NZ / NrxIV-GFP

Figure 6A

Ap-Gal4 / + ; UAS-Dcr2 , UAS-LacZ.NZ / UAS-LacZ.NZ

Ap-Gal4 / UAS-Kune^RNAi^ ; UAS-Dcr2 , UAS-LacZ.NZ / +

Figure 6B,C

Bx-Gal4 / + ; UAS-Dcr2 / + ; UAS-LacZ.NZ / +

Bx-Gal4 / + ; UAS-Dcr2 / UAS-Kune^RNAi^

Bx-Gal4 / + ; UAS-Dcr2 / UAS-EcR.A^W650A^

Figure 6D

Bx-Gal4 / + ; UAS-Dcr2 / + ; UAS-LacZ.NZ / +

Figure 6D

Bx-Gal4 / + ; UAS-Dcr2 / UAS-Cora^RNAi^

Figure 6D

Bx-Gal4 / + ; UAS-Dcr2 / UAS-EcR.A^W650A^

Figure 7A

Bx-Gal4 / + ; UAS-Dcr2 / + ; UAS-LacZ.NZ / +

Bx-Gal4 / + ; UAS-Dcr2 / UAS-Kune^RNAi^

Bx-Gal4 / + ; UAS-Dcr2 / + ; UAS-Eiger / +

Bx-Gal4 / + ; UAS-Dcr2 / UAS-Kune^RNAi^ ; UAS-Eiger / +

Figure S1A,C

Ap-Gal4 / + ; UAS-Dcr2 , UAS-LacZ.NZ / UAS-LacZ.NZ

Figure S1B,C

Ap-Gal4 / + ; UAS-Dcr2 , UAS-LacZ.NZ / UAS-Dilp8::3xFLAG

Figure S2A

Bx-Gal4 / + ; UAS-Dcr2 / + ; UAS-LacZ.NZ / +

Bx-Gal4 / + ; UAS-Dcr2 / UAS-Nrx^RNAi^

Bx-Gal4 / + ; UAS-Dcr2 / + ; UAS-Dilp8::3xFLAG / +

Bx-Gal4 / + ; UAS-Dcr2 / UAS-Nrx^RNAi^ ; UAS-Dilp8::3xFLAG / +

Figure S2B

Bx-Gal4 / + ; UAS-Dcr2 / + ; Dilp8^MI00727^ / Dilp8^MI00727^

Bx-Gal4 / + ; UAS-Dcr2 / UAS-Nrx^RNAi^ ; Dilp8^MI00727^ / Dilp8^MI00727^

Bx-Gal4 / + ; UAS-Dcr2 / + ; UAS-Dilp8::3xFLAG , Dilp8^MI00727^ / Dilp8^MI00727^

Bx-Gal4 / + ; UAS-Dcr2 / UAS-Nrx^RNAi^ ; UAS-Dilp8::3xFLAG , Dilp8^MI00727^ / Dilp8^MI00727^

Figure S3B

Ap-Gal4 / + ; UAS-Dcr2 , UAS-LacZ.NZ / UAS-LacZ.NZ

Ap-Gal4 / UAS-Kune^RNAi^ ; UAS-Dcr2 , UAS-LacZ.NZ / +

Figure S4

Ap-Gal4 / + ; UAS-Dcr2 , UAS-LacZ.NZ / UAS-LacZ.NZ

Ap-Gal4 / UAS-Kune^RNAi^ ; UAS-Dcr2 , UAS-LacZ.NZ / +

Figure S5

Ap-Gal4 / + ; UAS-Dcr2 , UAS-LacZ.NZ / UAS-LacZ.NZ

Ap-Gal4 / UAS-Kune^RNAi^ ; UAS-Dcr2 , UAS-LacZ.NZ / +

Figure S6

NrxIV-GFP / NrxIV-GFP

Figure S7

NrxIV-GFP / NrxIV-GFP

Figure S8B,C

Ap-Gal4 / UAS-Kune^RNAi^ ; UAS-Dcr2 , UAS-LacZ.NZ / NrxIV-GFP

Figure S8D,E

Ap-Gal4 / UAS-Nrx^RNAi^ ; UAS-Dcr2 , UAS-LacZ.NZ / NrxIV-GFP

Figure S8F,G

Ap-Gal4 / UAS-Cora^RNAi^ ; UAS-Dcr2 , UAS-LacZ.NZ / NrxIV-GFP

Figure S9B

Ap-Gal4 / + ; UAS-Dcr2 , UAS-LacZ.NZ / UAS-LacZ.NZ

Ap-Gal4 / UAS-Kune^RNAi^ ; UAS-Dcr2 , UAS-LacZ.NZ / +

Figure S10B

Bx-Gal4 / + ; UAS-Dcr2 / + ; UAS-LacZ.NZ / +

Figure S10C

Bx-Gal4 / + ; UAS-Dcr2 / UAS-Cora^RNAi^

Figure S10D

Bx-Gal4 / + ; UAS-Dcr2 / UAS-EcR.A^W650A^

Figure S11

Bx-Gal4 / + ; UAS-Dcr2 / + ; UAS-LacZ.NZ / +

Bx-Gal4 / + ; UAS-Dcr2 / UAS-Nrx^RNAi^

Bx-Gal4 / + ; UAS-Dcr2 / + ; UAS-Eiger / +

Bx-Gal4 / + ; UAS-Dcr2 / UAS-Nrx^RNAi^ ; UAS-Eiger / +

Figure S12

Bx-Gal4 / + ; UAS-Dcr2 / + ; UAS-Eiger / Dilp8^MI00727^

Bx-Gal4 / + ; UAS-Dcr2 / UAS-Kune^RNAi^ ; UAS-Eiger / Dilp8^MI00727^

Bx-Gal4 / + ; UAS-Dcr2 / UAS-Nrx^RNAi^ ; UAS-Eiger / Dilp8^MI00727^

**Supplementary Figures**

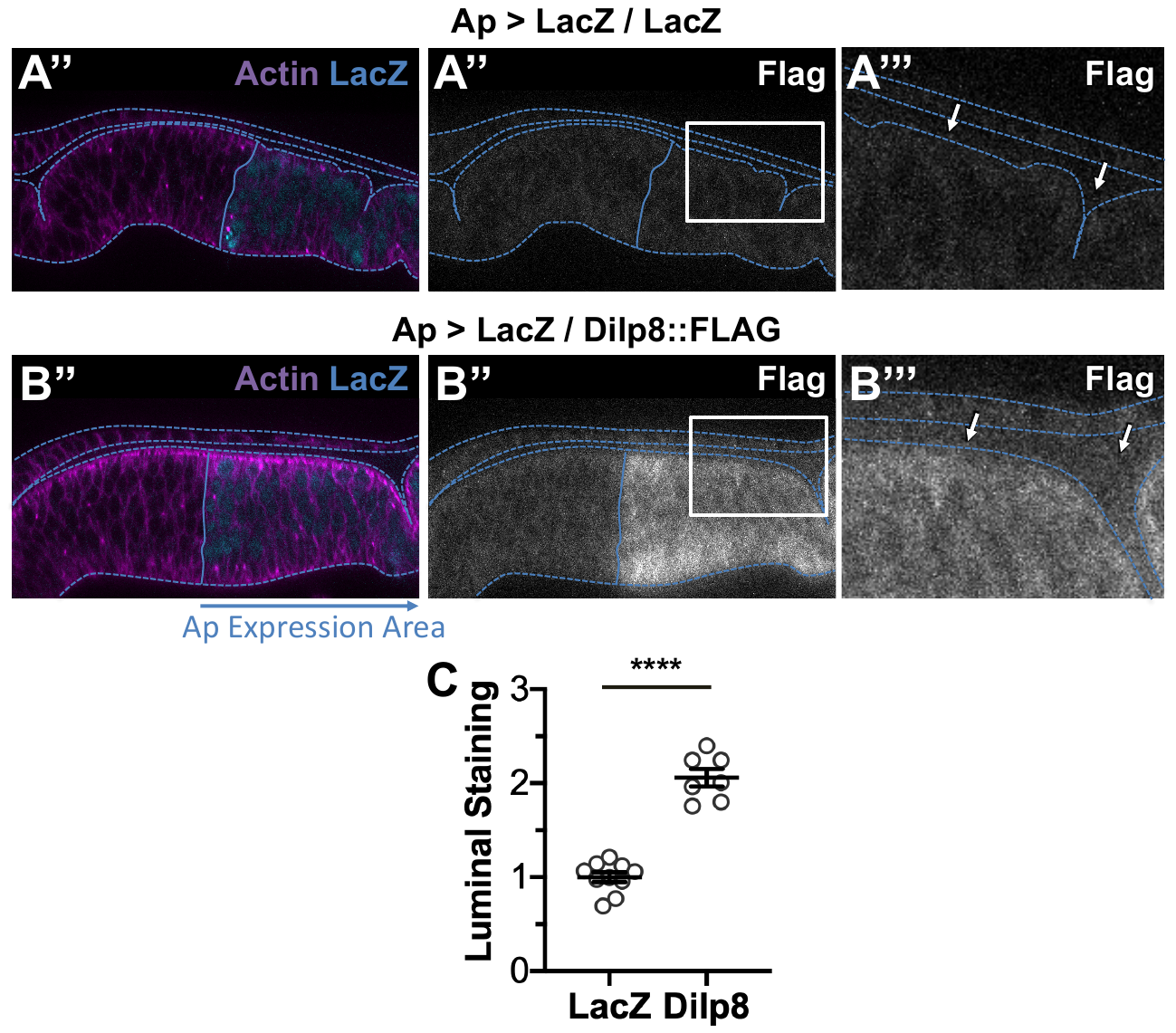

**Figure S1.** Dilp8 accumulates in the wing imaginal disc lumen. Ap-Gal4 was used to express (A) LacZ (wild type control) or (B) LacZ and Dilp8::FLAG in the dorsal region of wing imaginal discs. (A-B) Images are XZ cross-sections of wing imaginal discs in the pouch region of the disc. Dotted blue lines indicate disc area as defined by Actin (rhodamine phalloidin staining). Solid blue lines indicate the dorsal-ventral boundary, as defined by LacZ expression (β-Gal staining). Representative images are oriented dorsal on the right. (A) No FLAG is observed in *Ap > lacZ / lacZ* expressing discs either in (A’’) the expression region or (A’’’) the lumen. (B) FLAG is observed in *Ap > lacZ / dilp8::FLAG* expressing discs in the (B’’) expression region and (B’’’) the lumen. (C) Quantification of FLAG in the lumen, normalized to *Ap > LacZ* expression. Graph represent mean ± SEM, with individual points indicating values of single images. n = (LacZ) 10 and (Dilp8) 7 discs. **** p < 0.0001 by unpaired t-test.

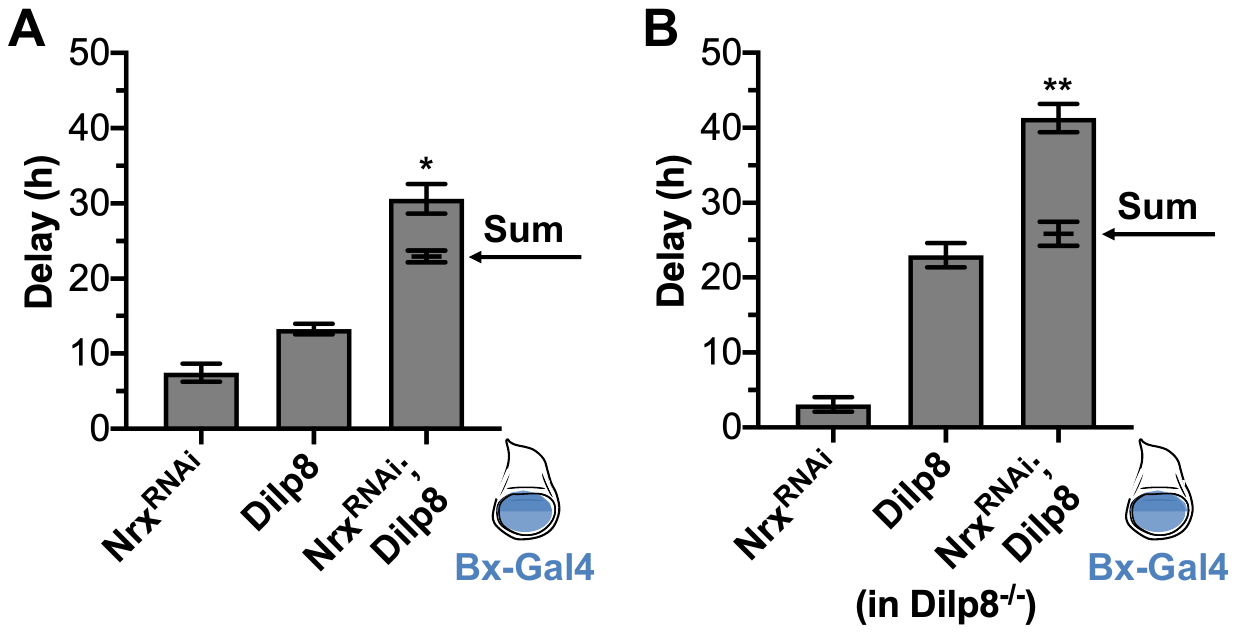

**Figure S2.** The epithelial barrier limits Dilp8 signaling. (A) Co-expression of *nrx^RNAi^* and Dilp8 induces synergistic delay. Ectopic expression of nrx^RNAi^, Dilp8, and co-expression of *nrx^RNAi^* and Dilp8 (*nrx^RNAi^*; Dilp8) induce developmental delay compared to LacZ controls when expressed in the wing imaginal disc under Bx-Gal4 (expression region in blue). The delay induced by co-expression of *nrx^RNAi^* and Dilp8 (*nrx^RNAi^*; Dilp8) is significantly more than the sum of the delay induced by *nrx^RNAi^* and Dilp8 expressed alone (sum indicated by arrow). (B) This trend holds true when endogenous Dilp8 is limited by expression in a Dilp8 hypomorphic background (Dilp8^MI00727^/Dilp8^MI00727^; (Garelli et al., 2012)). Data were collected from at least four independent experiments, bars represent mean ± SEM, * p < 0.05, ** p < 0.01 from one sample t-test comparing the additive value and observed delay.

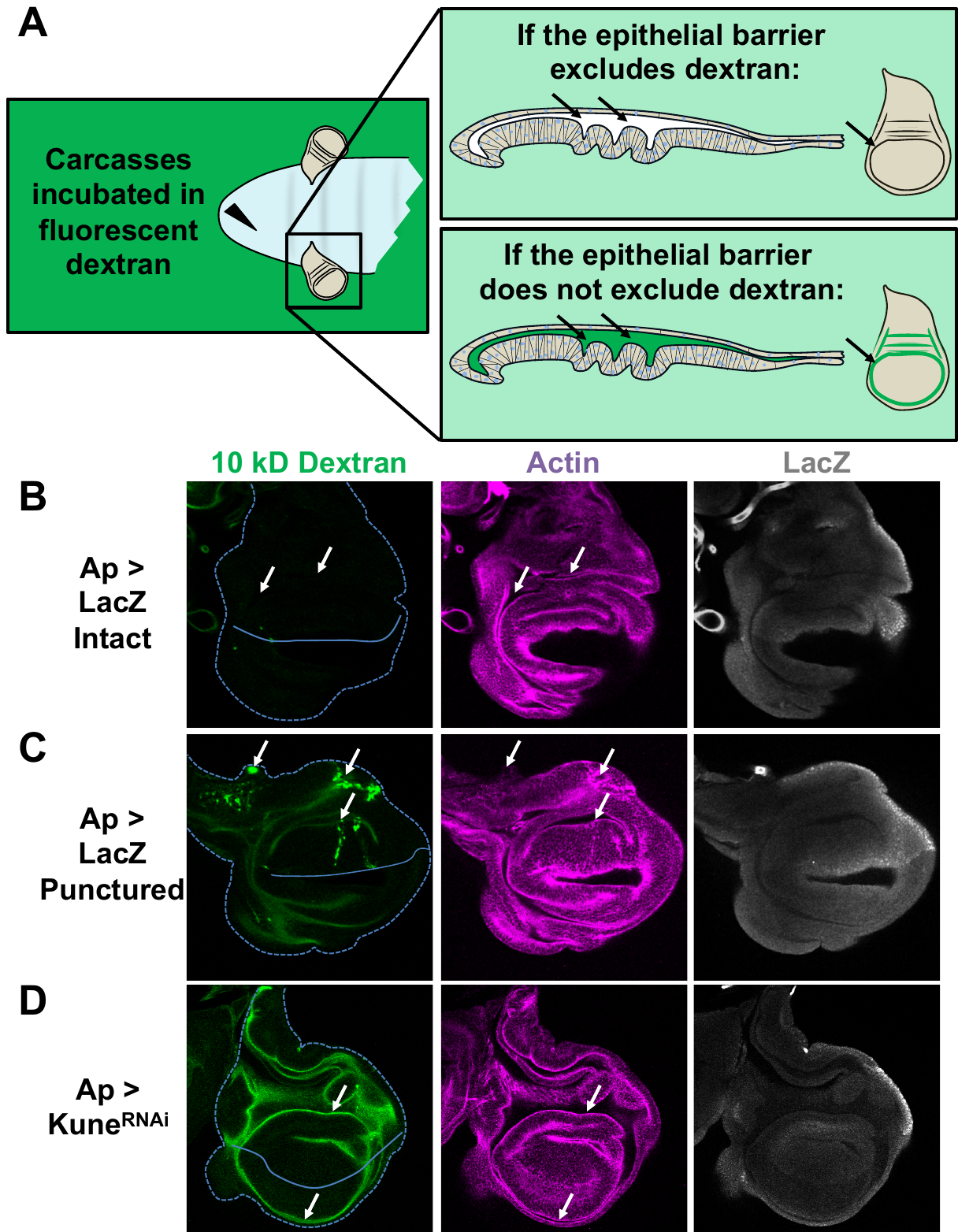

**Figure S3.** Explanation of the dextran assay for barrier function. (A) Carcasses are inverted and cleaned, then incubated in a fluorescence conjugated dextran for 30 minutes before fixation, the discs were then stained, mounted, and imaged (full description in methods). If the epithelial barrier excludes the dextran, no dextran should be observable in the lumen of the imaginal disc. If the epithelial barrier does not exclude the dextran or the tissue integrity is disrupted, dextran should be observable in the lumen. (B-D) Representative images after 10 kD fluorescein-conjugated dextran incubation (from the experiment quantified in Figure 2A). Ap-Gal4 was used to express LacZ (Ap > LacZ) or Kune^RNAi^ (Ap > Kune^RNAi^). Dextran is observed in discs punctured LacZ expressing discs (C) and Kune^RNAi^ expressing discs (D), but not in intact LacZ expressing discs (A). Disc area is indicated by the dashed line, as defined by Actin staining (rhodamine phalloidin). Area of expression is dorsal (oriented up) of the solid line, as defined by LacZ staining (b-Gal; note that LacZ expressing discs express two copies of LacZ while Kune^RNAi^ expressing discs have one copy so β-Gal staining is not comparable between images). Arrows indicate: (A) areas of the lumen with no distinguishable dextran fluorescence; (B) areas where damage during dissection (puncturing) has disrupted epithelial barrier integrity, allowing dextran to enter the cells and the imaginal disc lumen; and (C) luminal dextran is observed in both the dorsal lumen (Kune^RNAi^ expressing area, oriented up) and also in the ventral lumen (non-expressing area, oriented down), indicating that the lumen is contiguous. Images are single slices.

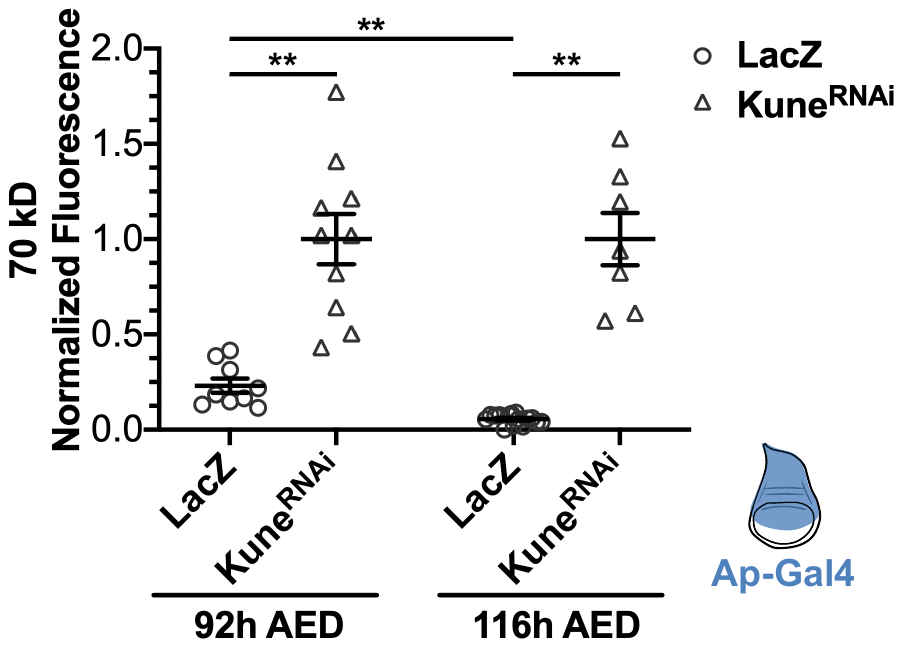

**Figure S4.** The epithelial barrier of wing imaginal discs grow more exclusionary to 70 kD fluorescent dextran during the third instar. The function of the epithelial barrier to exclude 70 kD Texas Red conjugated dextran was measured, as previously described, at 92h and 116h AED in wing imaginal discs expressing LacZ or Kune^RNAi^ by Ap-Gal4 (expression area diagramed in blue). Data are normalized to the mean luminal intensity of the Kune^RNAi^ expressing discs. Graph represents mean ± SEM, with individual points indicating values of single images. Left to right, n = 9, 10, 15, 7. ** p < 0.01 as calculated by Brown-Forsythe and Welch ANOVA with Dunnett’s T3 test for multiple comparisons.

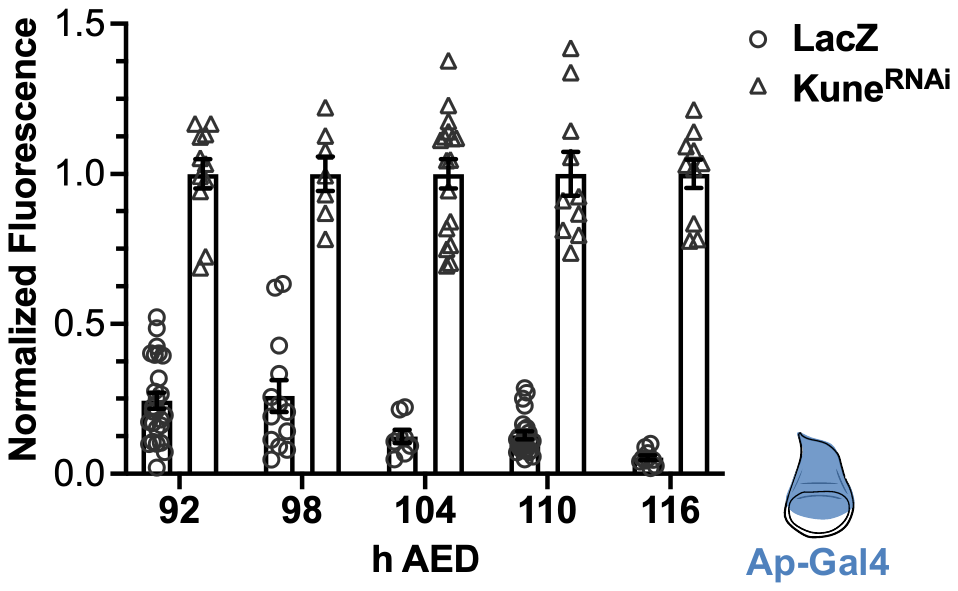

**Figure S5.** The epithelial barrier matures between 92h and 116h AED, becoming more restrictive to 10 kD fluorescein conjugated dextran. These are the complete data from Figure 2C, including data from Kune^RNAi^ expressing discs: the barrier function of wing imaginal discs expressing LacZ or Kune^RNAi^ by Ap-Gal4 was measured every 6 hours between 92h and 116h AED. Data indicate luminal intensity of intact LacZ expressing discs normalized to the mean luminal intensity of the Kune^RNAi^ expressing discs from the same timepoint. Graphs represent mean ± SEM, with individual points indicating values of single images. Significance between LacZ expressing discs at each timepoint is indicated in Figure 2C. Left to right, n = (92h AED) 26, 11, (98h AED) 13, 7, (104h AED) 12, 17, (110h AED) 25, 10, (116h AED) 10, 11.

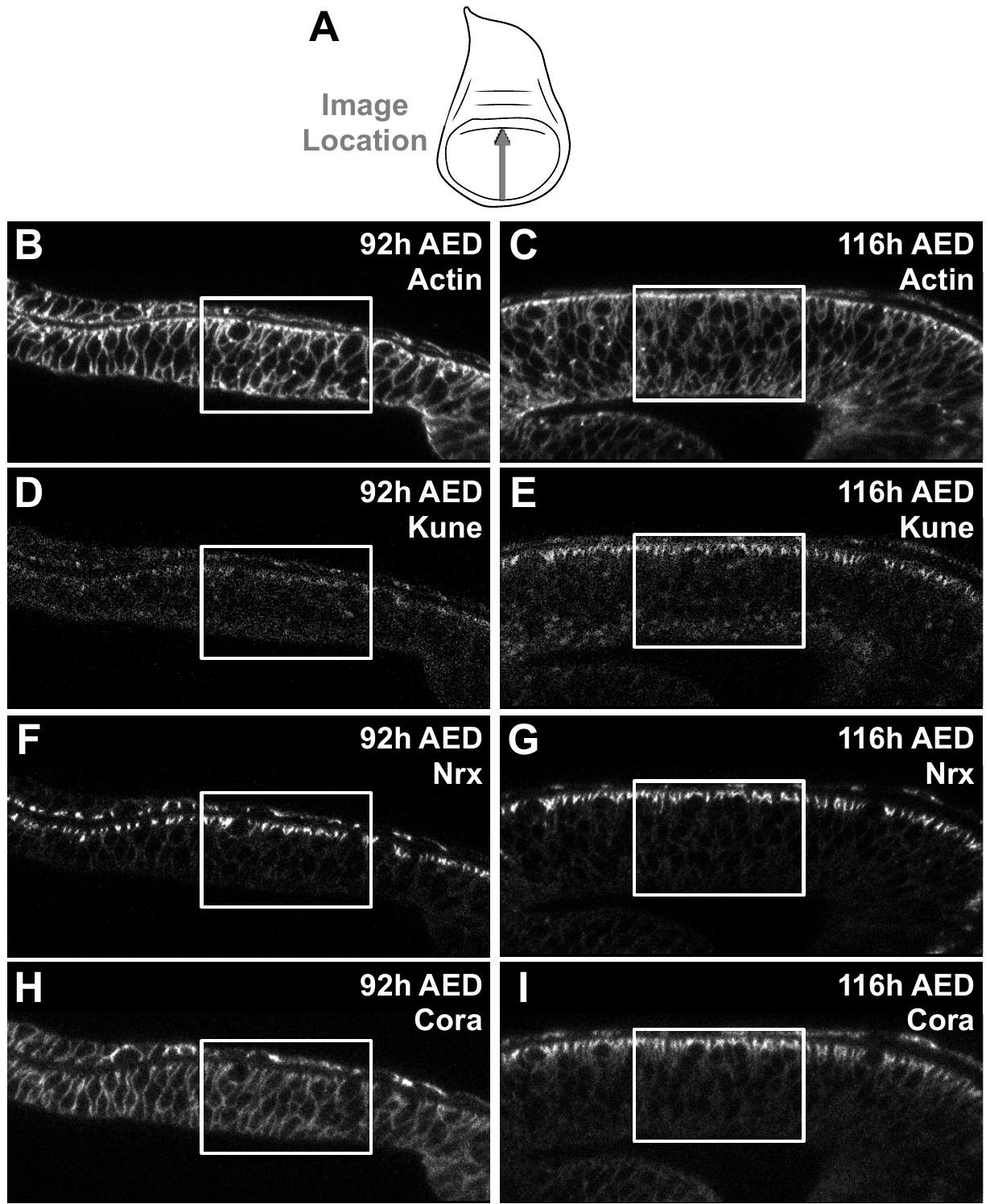

**Figure S6.** Complete images from Figure 3. (A) Approximate area and orientation of XZ image locations in the wing imaginal discs. (B-I) Representative XZ images at 92h and 116h AED, the region zoomed into in Figure 3 is indicated (white box). Images are: (B-C) Actin (rhodamine phalloidin; corresponds to Figure 3 B-C), (D-E) Kune (Anti-Kune; corresponds to Figure 3 E-F), (F-G) Nrx (Nrx-GFP; corresponds to Figure 3 G-H); and (H-I) Cora (Anti-Cora; corresponds to Figure 3 J-K).

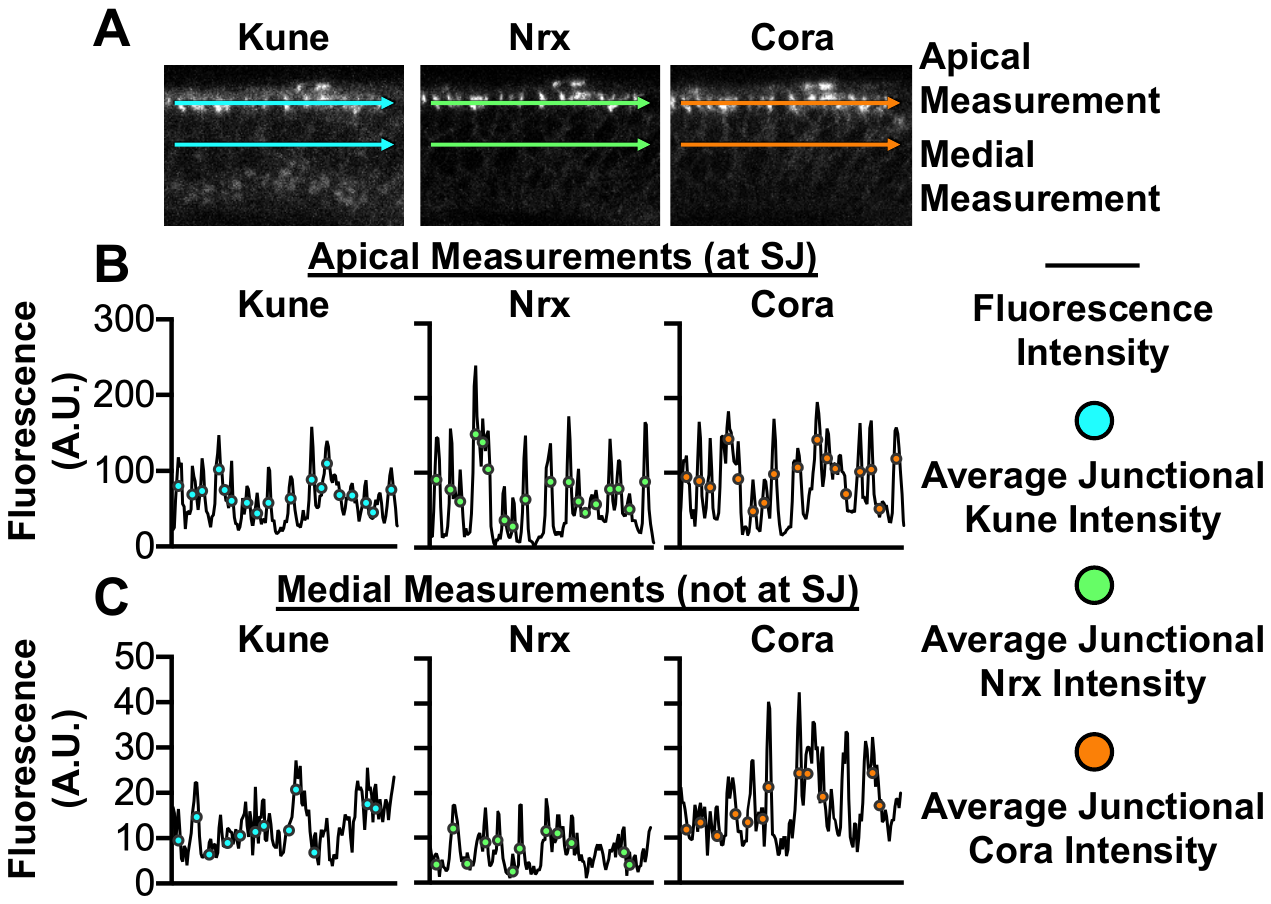

**Figure S7.** Method for junction quantification. (A) Lines were drawn apically bisecting the region of brightest septate junction (SJ) staining, and medially. (B-C) Fluorescence intensity across each line were measured with plot profile. Areas of the membrane (septate junction if apical) were identified as local maxima (peak) within a 7-pixel range, the 3 prior and following the pixel in questions. We adjusted these data in three steps, further described in Materials and Methods, to reduce false identification of a peak being localized at a membrane due to imaging or staining issues (eg. non-specific staining, image noise), especially with regards to Anti-Kune and Anti-Cora staining. The average junctional intensity was taken as the mean of the 7-pixel range.

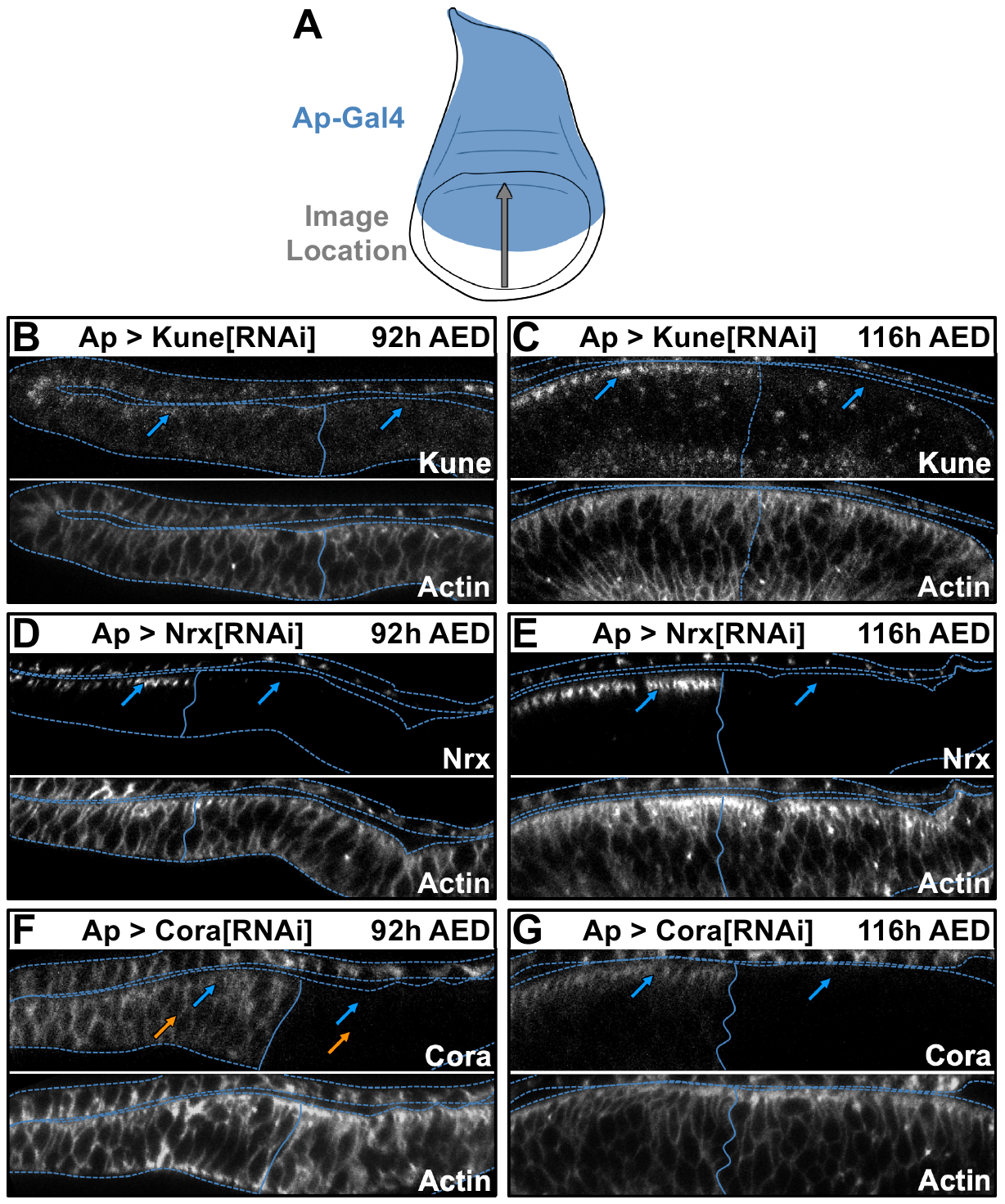

**Figure S8.** RNAi inhibition of septate junction components. (A) Images were taken spanning the dorsal-ventral boundary in the pouch region of wing imaginal discs (approximate area indicated by grey arrow), which includes tissue outside and inside the Ap-Gal4 expression region (blue). (B-G) Localization of septate junction components following RNAi expression and actin localization (defined by rhodamine phalloidin). Images are from the same discs as Figure 5. Dotted line represents tissue outline defined by actin staining. Solid line represents dorsal-ventral boundary, expression area (dorsal region) is on the right. Blue arrows indicate apical-lateral localization, orange arrows indicate medial-lateral localization. (B-C) Localization of Kune and Actin in *Ap > kune^RNAi^* expressing discs at (B) 92h and (C) 116h AED. Kune is depleted in the *kune^RNAi^* expression region at both times. Images are of the same discs as Figure 4A and 4B. (D-E) Localization of Nrx and Actin in *Ap > nrx^RNAi^* expressing discs at (D) 92h and (E) 116h AED. Nrx is depleted in the *nrx^RNAi^* expression region at both times. Images are of the same discs as Figure 4C and 4D. (F-G) Localization of Cora and Actin in *Ap > cora^RNAi^* expressing discs at (F) 92h and (G) 116h AED. Kune is depleted in the *cora^RNAi^* expression region at both times. Images are of the same discs as Figure 4E and 4F.

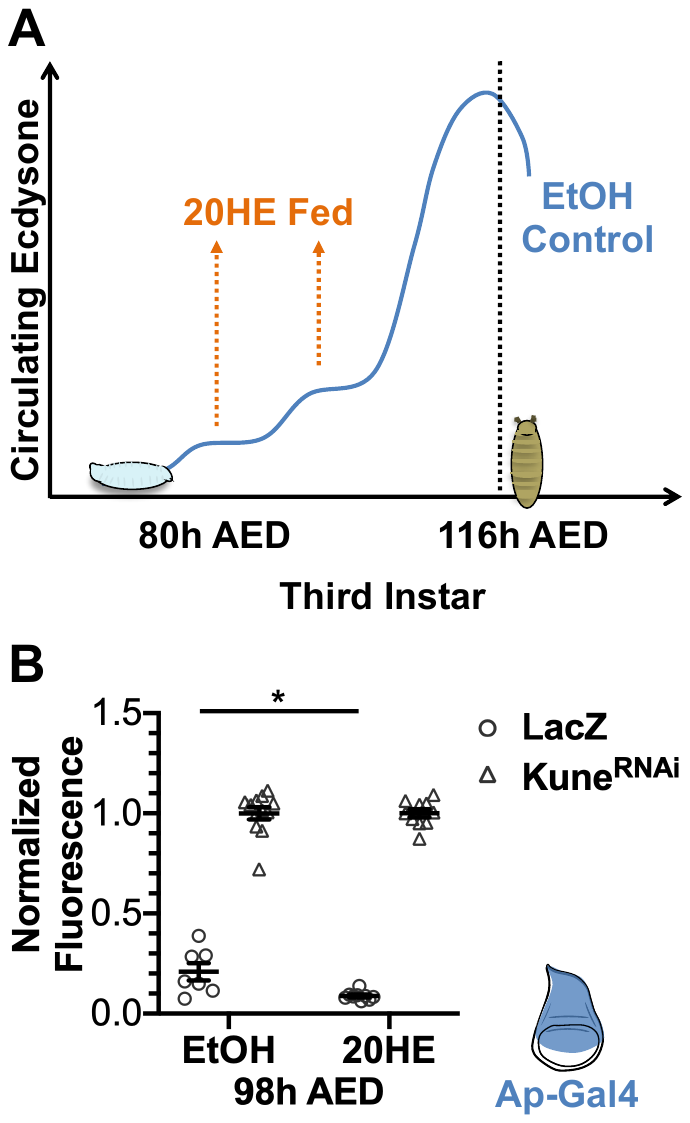

**Figure S9.** Ecdysone feeding induces barrier maturation early. (A) At 80h AED, larvae were switched to food containing 0.6 mg/mL 20-hydroxyecdysone (20HE) or ethanol control (EtOH). Feeding 0.6 mg/mL 20HE at 80h AED does not influence pupariation time, but does limit regeneration (Colombani et al., 2005; Jaszczak et al., 2015). (B) Wing imaginal disc barrier function at 98h AED of larvae fed the EtOH or 20HE food. Data are from larvae expressing *Ap > lacZ* (data in Figure 6A) or *Ap > kune^RNAi^.* Expression area indicated in blue. Graph represents mean ± SEM, with individual points indicating values of single images. Left to right, n = 7, 12, 9, 10. * p < 0.05 as calculated by unpaired t-test with Welch’s correction.

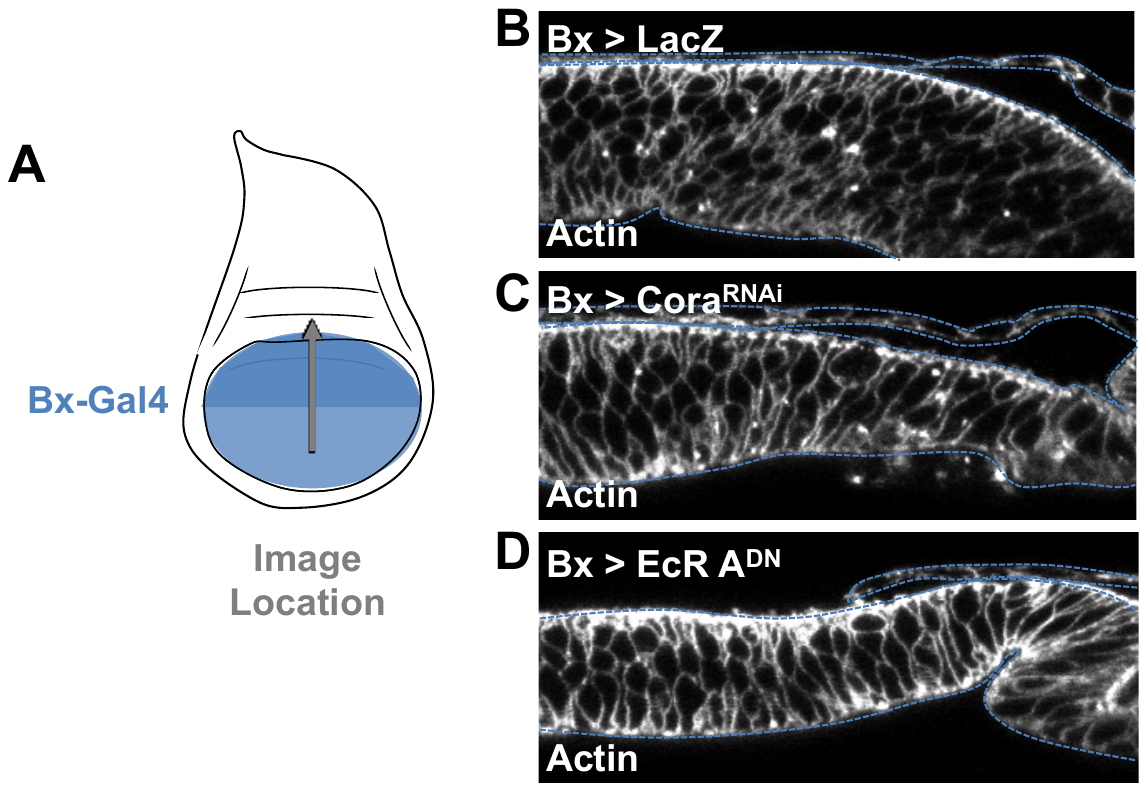

**Figure S10.** Actin stain from discs in Figure 6. (A) Bx-Gal4 expression area (blue) and approximate image location (grey arrow) for B-D. (B-D) Localization of Actin (rhodamine phalloidin) in (B) *Bx > lacZ* (wild type control), (C) *Bx > kune^RNAi^*, and (D) *Bx > EcR A^DN^* at 116h AED. Dotted lines indicate tissue outline. Tissues are oriented with dorsal on the right. Images are from the same discs as Figure 6D-F.

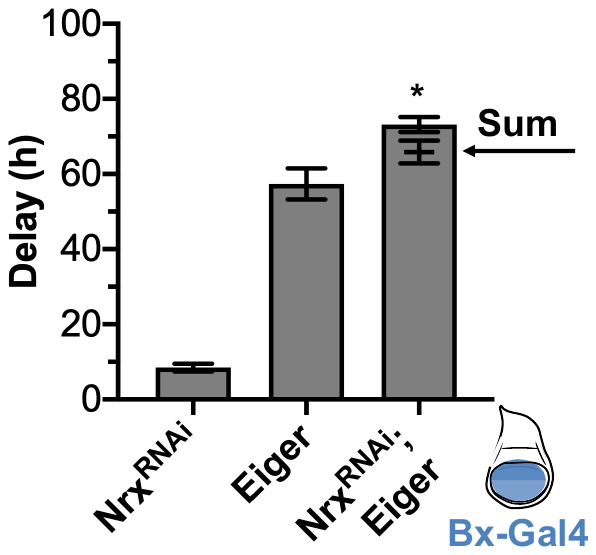

**Figure S11.** Nrx limits Eiger-induced delay. Co-expression of *nrx^RNAi^* and Eiger produces synergistic delay. Ectopic expression of *nrx^RNAi^*, Eiger, and co-expression of *nrx^RNAi^* and Eiger (*nrx^RNAi^*; Eiger) induce developmental delay compared to LacZ controls when expressed in the wing imaginal disc under Bx-Gal4 (expression region in blue). The delay induced by co-expression of *nrx^RNAi^* and Eiger (*nrx^RNAi^*; Eiger) is significantly more than the sum of the delay induced by *nrx^RNAi^* and Eiger expressed alone (sum indicated by arrow. Data were collected from at least four independent experiments, bars represent mean ± SEM, * p < 0.01 from one sample t-test comparing the additive value and observed delay.

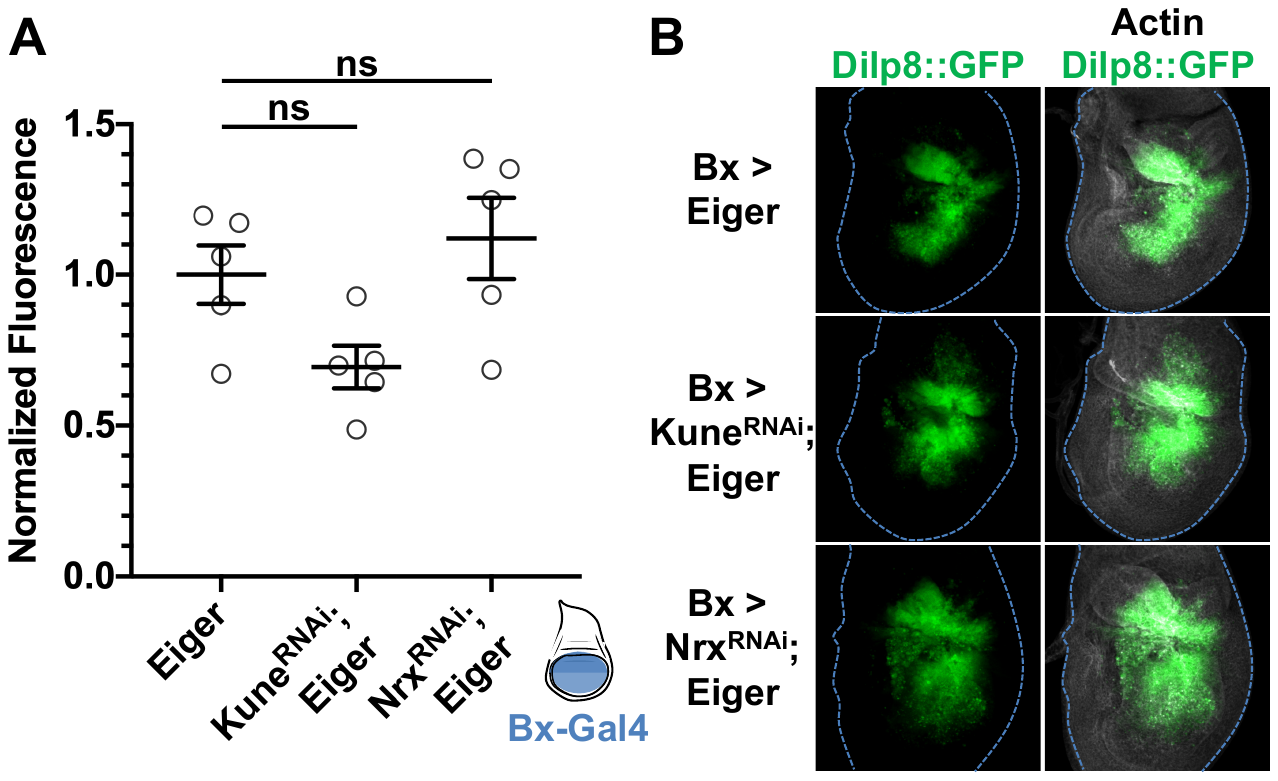

**Figure S12.** Coexpression of Eiger and RNAi against septate junction components does not significantly alter measured Dilp8 expression. Bx-Gal4 (expression area in blue) was used to express Eiger alone, with *kune^RNAi^*, or with *nrx^RNAi^* expressing a transcriptional reporter for Dilp8 (Dilp8^MI00727^/+; Garelli et al., 2012). (A) Sum GFP intensity was measured and the data were normalized to Eiger alone. (B) Representative images. Discs were collected at 104 hAED. Images represent sum-projected stacks of 5 images. Actin was stained for with rhodamine phalloidin. (A) Graph represents mean ± SEM, with individual points indicating values of single images, n = 5 images in each condition; ns indicates p > 0.05 by one-way ANOVA with Tukey test for multiple comparisons.

**Supplemental References**

Colombani, J., Bianchini, L., Layalle, S., Pondeville, E., Dauphin-Villemant, C., Antoniewski, C., Carré, C., Noselli, S. and Léopold, P. (2005). Antagonistic actions of ecdysone and insulins determine final size in Drosophila. *Science* 310, 667–670.

Garelli, a., Gontijo, a. M., Miguela, V., Caparros, E. and Dominguez, M. (2012). Imaginal Discs Secrete Insulin-Like Peptide 8 to Mediate Plasticity of Growth and Maturation. *Science* 336, 579–582.

Jaszczak, J. S., Wolpe, J. B., Dao, A. Q. and Halme, A. (2015). Nitric oxide synthase regulates growth coordination during Drosophila melanogaster imaginal disc regeneration. *Genetics* 200, 1219–1228.
